# Restricted constrictors: Space use and habitat selection of native Burmese pythons in Northeast Thailand

**DOI:** 10.1101/2020.09.17.302661

**Authors:** Samantha Nicole Smith, Max Dolton Jones, Benjamin Michael Marshall, Surachit Waengsothorn, George A. Gale, Colin Thomas Strine

## Abstract

Animal movement and resource use are tightly linked. Investigating these links to understand how animals utilize space and select habitats is especially relevant in areas that have been affected by habitat fragmentation and agricultural conversion. We set out to explore the space use and habitat selection of Burmese pythons (*Python bivittatus*) in a patchy land use matrix dominated by agricultural crops and human settlements. We used radio telemetry to record daily locations of seven Burmese pythons over the course of our study period of approximately 22 months. We created dynamic Brownian Bridge Movement Models (dBBMMs) for all individuals, using occurrence distributions to estimate extent of movements and motion variance to reveal temporal patterns. Then we used integrated step selection functions to determine whether individual movements were associated with particular landscape features (aquatic agriculture, forest, roads, settlements, terrestrial agriculture, water), and whether there were consistent associations at the population level. Our dBBMM estimates suggested that Burmese pythons made use of small areas (98.97 ± 35.42 ha), with low mean individual motion variance characterized by infrequent moves and long periods at a single location. At both the individual and population level, Burmese pythons in the agricultural matrix were associated with aquatic environments. Only one individual showed a strong avoidance for human settlements which is troublesome from a human-wildlife conflict angle, especially as Burmese pythons have been observed entering human settlements and consuming livestock in our study site. Our study is one of the first to contribute to the knowledge of Burmese python ecology in their native range as the majority of studies have focused on invasive populations.

## Introduction

Studying and understanding animal movement allows scientists to gain insight into animal behavior (Gurarie et al., 2009), revealing information about species’ natural and life histories such as migratory or movement patterns, breeding grounds and seasons, foraging strategies, and habitat selection (Block et al., 2011; Dulau et al., 2017; Glaudas & Alexander, 2016; Viana et al., 2018). Animal movement is directly connected to resource use such as habitat selection (Moorter et al., 2016). As humans continue to encroach on suitable habitats, understanding relationships between animal movement and habitat selection can assist in the development of conservation-based actions.

Tropical regions disproportionately undergo rapid deforestation and land-use changes compared to non-tropical forests (Austin et al., 2017; Hansen et al., 2013; Laurance et al., 2014); this rate is especially alarming as many of the world’s biodiversity hotspots occur in tropical regions (Mittermeier et al., 1998; Myers et al., 2000). Southeast Asia in particular is vulnerable to biodiversity loss and suffers from the highest deforestation rate of any region (Hughes, 2017; Sodhi et al., 2004), with land being transformed to accommodate growing populations, subsequent urban sprawl and agricultural conversion (Laurance, 2007; Laurance et al., 2014; Schneider et al., 2015).

Of species affected by habitat encroachment, land use-change and deforestation, reptiles are among some of the most heavily impacted and are experiencing population declines worldwide (Gibbons et al., 2000; Todd et al., 2010). Gibbons et al. (2000) list habitat loss, introduced invasive species, pollution, and climate change as some of the factors contributing to population declines. Reptiles are also commonly collected from the wild and traded in the exotic pet trade, which has been considered the second largest threat to the conservation of reptiles (Böhm et al., 2013). Regardless of widespread population declines, reptiles are generally understudied (Böhm et al., 2013; Gibbons et al., 2000) and therefore vast knowledge gaps about their ecology exist. Of the 45% (4648) of described reptile species assessed by the International Union for Conservation of Nature (IUCN), 19% (867) of these species are classified as data deficient (DD) (Tingley et al., 2016). The paucity of ecological studies on reptiles often means that even seemingly “well-known” species have very little information available about their population dynamics, distributions, resource use and fecundity. These substantial knowledge gaps about reptiles, especially in regions undergoing rapid transformation, highlight the importance of investigating aspects of their natural and life histories.

Habitat fragmentation can affect species differently, potentially putting taxa that occupy a more specialized ecological niche at a higher risk of extinction (Böhm et al., 2016). In contrast, species that are considered to be more generalist may be able withstand a changing landscape due to their ecological traits, or by making behavioral changes (Ditchkoff et al., 2006; McKinney, 2006; Shamoon et al., 2018). Carpet pythons (*Morelia spilota*), for example are able to survive in suburban areas in Australia by sheltering in vegetation thickets or even in the roofs of domestic residences and eating non-native species such as rats, pets, and livestock (Shine & Fitzgerald, 1996). While generalist species may be more resilient to habitat changes, their resilience and associated affinity to live in close proximity to humans often leads to human-wildlife conflict (Bateman & Fleming, 2012; Charles & Linklater, 2013; Soulsbury & White, 2015). Understanding how species living in modified areas move through and use the space in such areas can help us to better understand their ecology as well as develop conservation and human-wildlife conflict mitigation strategies.

The Burmese python (*Python bivittatus* KUHL 1820: 94) is a large, nonvenomous, constricting snake and is a known initiator of snake-human conflict. Burmese pythons are widely distributed throughout Southeast Asia and occur in Bangladesh, Myanmar, Laos, in Thailand north of the Isthmus of Kra, Cambodia, Vietnam, throughout Southern China and in disjunct populations in Northern and Northeastern India and Southern Nepal (Barker & Barker, 2008). In their native range, Burmese pythons use a wide range of habitat types such as tropical lowlands, grasslands, forests and within areas modified for human use (Barker & Barker, 2008; Cota, 2010; Rahman et al., 2014). Pythons occupy a broad ecological niche and consume a wide variety of prey items including birds, mammals and sometimes other reptiles (Bhupathy et al., 2014; Dorcas et al., 2012; Dove et al., 2011; Shine et al., 1998; Slip & Shine, 1988).

The IUCN lists Burmese pythons as Vulnerable (Stuart et al., 2019), due to harvesting for traditional medicine, the pet and skin trade, and habitat degradation (Stuart et al., 2019). Burmese pythons will enter human settlements and consume livestock, resulting in substantial human-wildlife conflict (Goodyear 1994; Rahman et al., 2014; You et al., 2013). Instances of human-wildlife conflict could potentially be a contributing factor to Burmese python population declines as persecution of snakes often occurs in conflict situations (Ceríaco, 2012; Miranda et al., 2016; Nóbrega Alves et al., 2012). For example, in a study investigating interactions between humans and anacondas in South America using internet videos, 52 (∼16%) of 330 videos assessed showed anacondas being killed as a result of these interactions (Miranda et al., 2016). Conflict between pythons and humans could perpetuate hostile attitudes towards snakes, as pythons can act as pest species via livestock predation.

Nearly all information on Burmese python ecology comes from their invasive range in the Florida Everglades, USA to determine their detrimental impact to native ecosystems (Dorcas et al., 2012; Dove et al., 2011; Hart et al., 2015; Hoyer et al., 2017; McCleery et al., 2015; Orzechowski et al., 2019; Willson, 2017). Only two studies have published results on native, free-ranging Burmese python movements –one tracked a single study animal for only 24 days in Hong Kong (Goodyear, 1994), the other tracked two males and two females for un-specified durations in Bangladesh (Rahman et al., 2014)– neither presenting sufficient information on Burmese python ecology, highlighting the need for further investigation. We currently have a very limited understanding on the space use of Burmese pythons, including any habitat preferences, importance of landscape features and resilience to human-mediated disturbances. To address this dearth of knowledge, we set out to (1) quantify space use (occurrence distributions during our study) and movements of adult Burmese pythons in a patchy land-use matrix and (2) identify how Burmese python movements are associated with certain habitat features (i.e., distance to aquatic agriculture, forest, roads, settlements, terrestrial agriculture, water).

## Methods

### Study site

We captured and tracked all study animals within the Sakaerat Biosphere Reserve in Nakhon Ratchasima Province, Thailand (14.44-14.55° N, 101.88-101.95° E) from 2018-09-29 to 2020-07-22 (Figure 1). The reserve is made up of a protected core area of 8,000 ha that includes several different forest types; dry evergreen forest (60%), dry dipterocarp forest (18%), and small patches of reforested area, grasslands, and bamboo forest making up the remaining area (22%). The core area is regularly patrolled by rangers employed by the Sakaerat Environmental Research Station (SERS) in attempt to prevent poaching of wildlife within the protected area. Just outside the protected area, there is an unprotected buffer zone and transitional zone that together make up 36,000 ha. The buffer zone encompasses stretches of unprotected or otherwise disturbed forests with patches of plantation forest regrowth scattered within. The transitional zone is mostly land used for agriculture and human settlements (Figure 2). Agricultural crops produced in the biosphere’s transitional zone include mainly rice paddy, cassava, sugarcane and corn. Climatic conditions in our study site can be generalized into three seasons; hot (mean: 33.8 ± standard error [SE] 2.8°C; 2.5 ± 7.9 mm rainfall), wet (29.9 ± 2.2°C; 5.9 ± 11.1 mm rainfall) and dry (29.0 ± 3.5°C; 0.2 ± 0.8 mm rainfall; Marshall et al., 2020).

**Figure 1.**
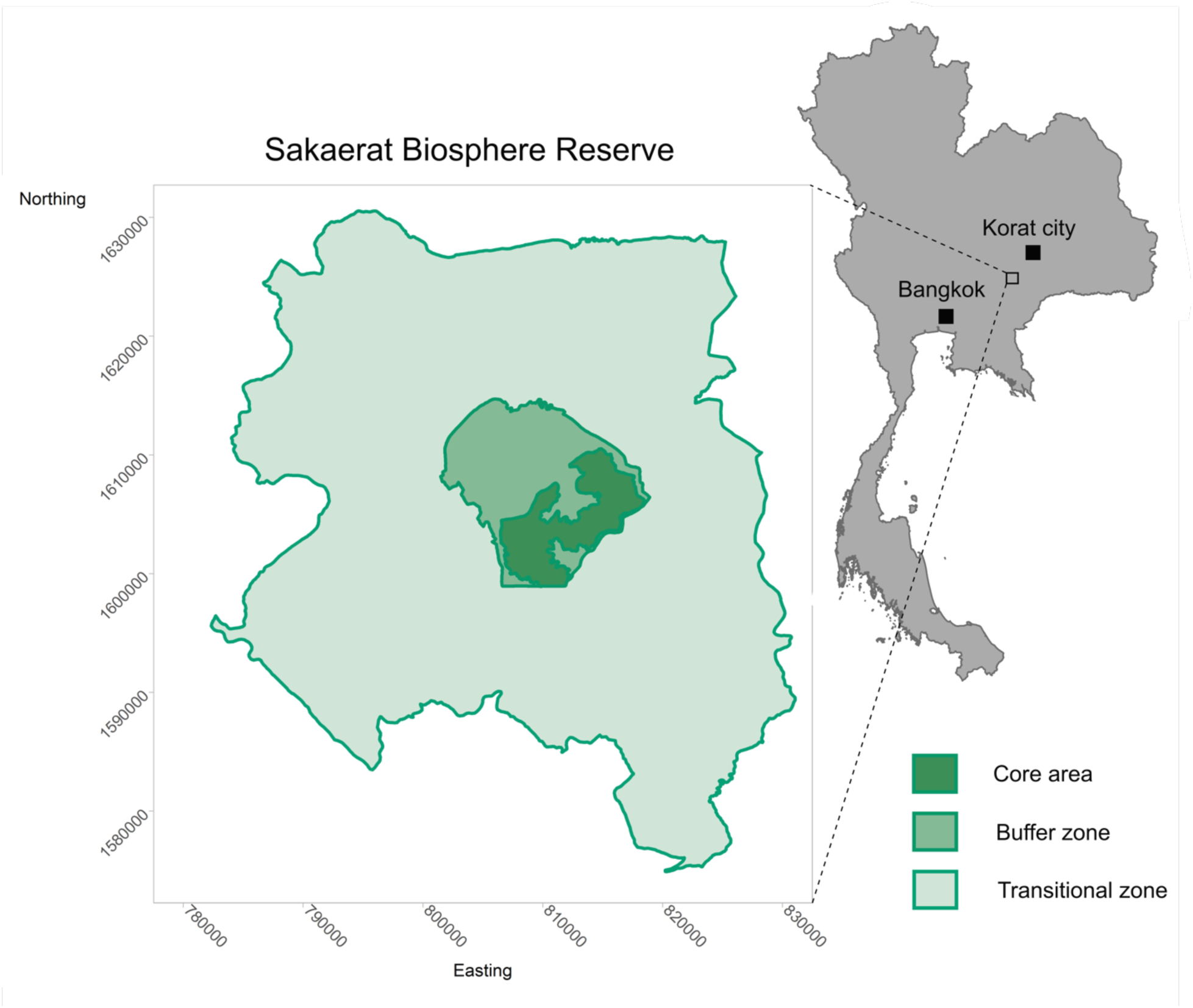
A map of the Sakaerat Biosphere Reserve with an inset map of Thailand highlighting the study site location in relation to Bangkok and Korat city.

**Figure 2.**
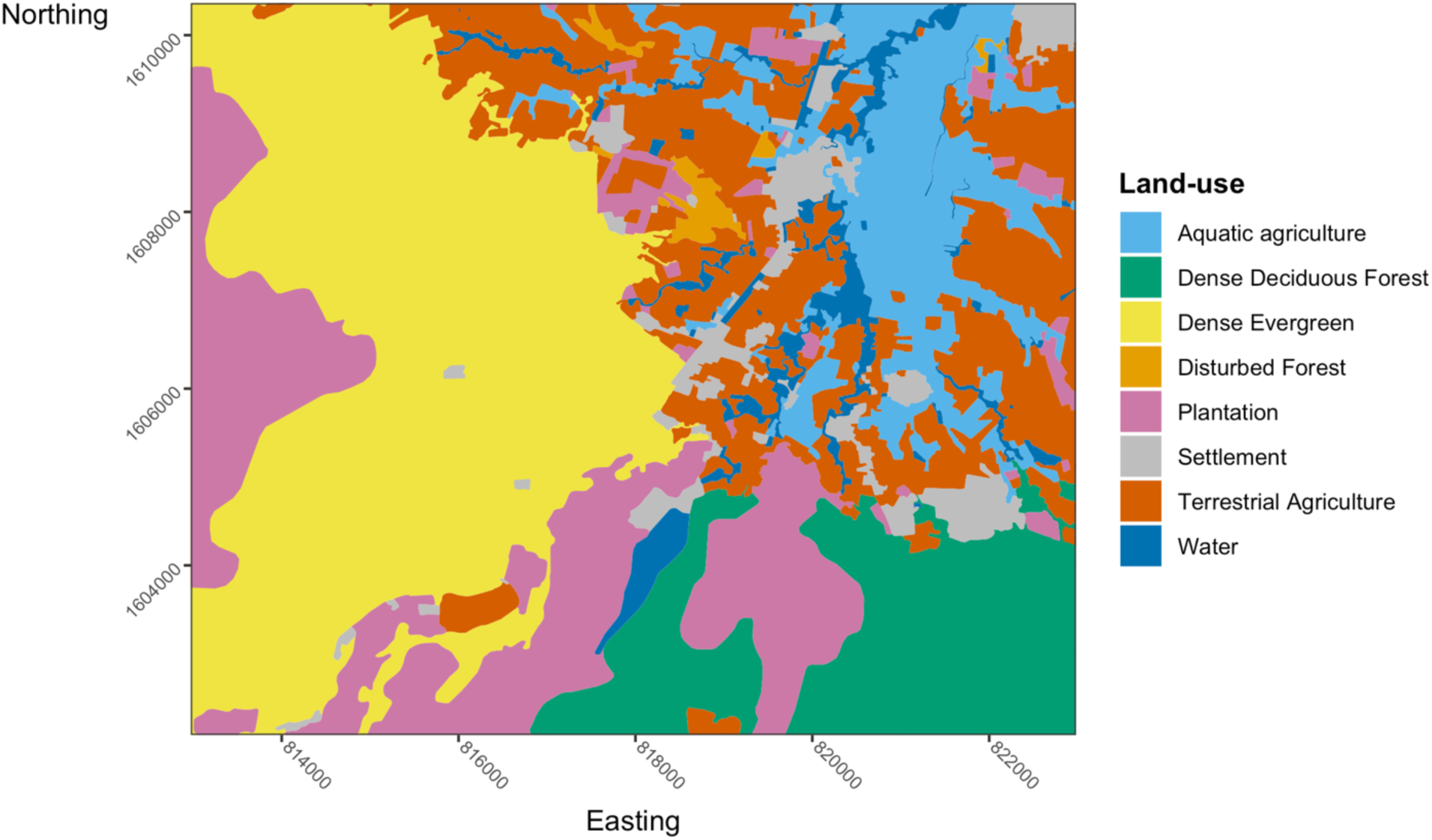
Land-use types spanning the area of which all study animals were tracked.

### Study animal capture, processing and implantation

We captured all study animals opportunistically, through visual encounter surveys, or via notification from residents in the surrounding villages (Supplementary Table 1). Once we captured an individual, we brought the snake back to our field laboratory at SERS to collect biometric data. We gave all snakes a unique ID using the first two letters of the genus and species and a number based off of the capture sequence (PYBI###). In order to successfully and accurately collect measurements, we used isoflurane, an inhalant anesthetic to temporarily subdue the snake. Once the snake was fully anesthetized, we collected biometric measurements such as snout to vent length (SVL; mm), tail length (TL; mm) and mass (g) (Table 1). We determined the sex of each snake via cloacal probing (Schaefer, 1934). In attempt to strengthen our statistical inferences and avoid sub-setting our sample size, we aimed to restrict transmitter-implantation to adult females. Using approximate estimates cited by Reed and Rodda (2009) we considered all female Burmese python with a total length of 2500 mm to be adults and therefore suitable for radio-tracking. All individuals acceptable for radio-tracking (i.e., females, 2500 mm or over, and captured with 15km of our study site) were surgically implanted with VHF radio transmitters (Holohil AI-2T or SI-2T, Holohil Inc., Ontario, Canada). All surgeries were performed by a licensed veterinarian from the Nakhon Ratchasima Zoo. We had to wait for the veterinarian to be available to perform surgeries meaning snakes were not implanted immediately after processing (Supplementary Table 1). Surgeries followed a modified methodology introduced by Reinert and Cundall (1982) with the addition of isoflurane. Following surgery, snakes were monitored and then released no more than 24 hours after implantation to provide maximum opportunity for natural thermoregulation during the healing process. We released snakes at their initial capture site; however, in instances where snakes were captured from local residences, we were not able to release them at the direct capture location. In these cases, we released snakes in areas that were away from other residences and had ample vegetation and cover. We released snakes an average of 149.12 m ± SE 49.49 m (range = 0 — 393m) from their initial capture site (Supplementary Table 1). We began data collection on implanted snakes the day following their release; as no obvious subsequent behavioral changes were observed from initially implanted and acclimated snakes, we included all tracking data in our analyses.

**Table 1.** ID and Biometric measurements of all radio tracked Burmese pythons

| ID | SVL (mm) | TL (mm) | Mass (g) | Sex |
| --- | --- | --- | --- | --- |
| PYBI021 | 2744 | 330 | 11240 | Female |
| PYBI022 | 2304 | 318 | 5040 | Female |
| PYBI028 | 2314 | 352 | 7785 | Male |
| PYBI029 | 2423 | 300 | 8660 | Female |
| PYBI033 | 2214 | 296 | 6780 | Female |
| PYBI055 | 3085 | 401 | 19485 | Female |
| PYBI060 | 2490 | 352 | 6965 | Female |

SVL: Snout to vent length, TL: tail length

### Radio-tracking

We radio-tracked study animals between daylight hours (06:00-18:30) one time per day. We had therefore planned for a number of datapoints equal to the number of days an individual was tracked; however, there were occasional instances that we could not track study animals due to staff limitations, equipment malfunction, not being able to locate the animal, inclement weather or the animal had been brought in to be re-implanted with a new transmitter. On average, snakes had 17 ± 7 missing data points throughout the study. Despite these instances, we maintained a mean tracking time lag of 25.43 ± 0.32 hours (Supplementary Figure 1). In order to increase the location accuracy, we homed-in on tracked snakes to pinpoint their location. In the event that we could not directly approach the snake (i.e., snake was in water body, terrain did not allow for us to approach) we used three-point triangulations from as close as possible to estimate the snake’s location. Upon locating the snake, we recorded Universal Transverse Mercator (UTM WGS-84) points using hand-held GPS units (Garmin 64s) and GPS accuracy (m). To decrease disturbance, once we located the snake, we would retreat at least 5 m before recording coordinates using our data collection forms.

### Data collection, management and software

We collected and recorded all radio-tracking data in the field, using the Epicollect 5 smartphone (Android and iOS) application and then directly uploaded to the Epicollect 5 cloud database. We downloaded and reviewed data monthly to double check for inconsistencies such as missing digits in GPS points recorded in the field, which we checked against points directly marked on handheld GPS units.

We conducted all data analysis and visualization in R v.3.6.3 (R Core Team, 2020) using RStudio v.1.2.1335 (RStudio Team, 2019). We prepared data for analyses using packages *dplyr v*. 1.0.2 (Wickham et al., 2020) *data*.*table* v.1.13.0 (Dowle & Srinivasan, 2020) *reshape2* v.1.4.4 (Wickham, 2007), *readr* v.1.3.1 (Wickham et al., 2018), *lubridate* v.1.7.9 (Grolemund & Wickham, 2011), and *stringr* v.1.4.0 (Wickham, 2019). We calculated sample means and standard error (mean ± SE) using package *pracma* v.2.2.9 (Borchers, 2019). We used packages *rgdal* v.1.5.16 (R. Bivand et al., 2020), *raster* v.3.3.13 (Hijmans, 2020) and *sp* v.1.4.2 (R. S. Bivand et al., 2013) to work with rasters and shapefiles. For plot visualization we used *ggplot2* v.3.3.2 (Wickham, 2016), *scales* v.1.1.1 (Wickham & Seidel, 2020), *ggthemes* v.4.2.0 (Arnold, 2019), *ggspatial* v.1.1.4 (Dunnington, 2018) and *cowplot* v.1.0.0 (Wilke, 2019).

### Space use/occurrence distributions analysis

We quantified space use or occurrence distributions using dynamic Brownian Bridge Movement Models (dBBMMs). We elected to use dBBMMs to quantify space use over more traditional methods such as minimum convex polygons (MCPs) and kernel density estimates (KDEs) due to the fact that these traditional methods tend to greatly overestimate or underestimate space use during the study period (Silva et al., 2018, Silva et al., 2020). In contrast, dBBMMs create a more refined visualization of animal movement by taking into account the order of animal relocations and the duration of time spent at each location (Kranstauber et al., 2012). The dBBMMs require window size and margin size input to estimate an animal’s movement capacity and detecting changes in movement based on behavioral states. The window and margin size used in dBBMMs should be biologically relevant to the study species and sampling regime (Kranstauber et al., 2012). To set our window and margin size we followed methodology similar to Marshall et al. (2020). Given that tracked Burmese pythons would often remain stationary for long periods of time (∼2 weeks), we selected a window size of 15 datapoints. For the margin size, we selected a margin of 3 datapoints (both window and margin size must be odd) as we were able to detect behavioral changes (stationary vs. active) over the course of approximately 2 data points.

In addition to space use estimates of individuals during the study period, dBBMMs also provide motion variance estimates for tracked individuals that can reveal changes in movement. We explored seasonal patterns in motion variance for three of our snakes that were concurrently tracked for a year or more and through all three seasons (PYBI021, PYBI022 and PYBI029). Marshall et al. (2020). determined date ranges and environmental thresholds via cluster analysis for the three seasons that occur in our study site which we used to determine our seasonal delineations.

We produced telemetered Burmese python occurrence distributions and motion variance with dBBMMs using the *move* package v.4.0.2 (Kranstauber et al., 2020). To extract contours from the occurrence distributions and calculate areas, we used the packages *adehabitatHR* v.0.4.18 (Calenge, 2006) and *rgeos* v.0.5.3 (R. Bivand & Rundel, 2020).

Within their utilization distributions, we also investigated individual site fidelity using recursive analysis in package *recurse* v.1.1.2 (Bracis et al., 2018). For each individual, we took the mean GPS error to determine the radius for sites to be used in the analysis. Revisits occurred when an individual moved away from a location and then returned at any point after 24 hours had elapsed. We further report on the residence times (h) individuals spent at shelter sites locations.

### Integrated step selection analysis

To investigate Burmese python habitat selection, we used integrated step selection functions (ISSFs) on both an individual and population level (our entire sample of pythons using agricultural systems). For individual selection, we created integrated step selection functions using package *amt* (Signer et al., 2018). We used distance to various habitat features to determine selection versus avoidance on an individual basis. To create ISSFs, we modified code from Marshall et al. (2020b). We used a land use shapefile from the Thai Land Development Department (2017) which we separated into several categories of land use to create raster layers. We converted raster layers into layers with continuous values by taking the Euclidean distances to habitat features of interest (i.e., water bodies, forest, terrestrial agriculture, human settlements, roads, and aquatic agriculture). We grouped water bodies (ponds, reservoirs, and irrigation canals) with semi-natural areas, which typically consist of overgrown vegetation bordering water bodies as the two habitat features were highly correlated with each other. To avoid zero-inflation of distances to feature values and produce more intuitive effect directions in the models, we inverted all our raster layers. Global positioning system (GPS) trackers can provide high temporal resolution movement datasets; high temporal resolution data and the high computational costs of ISSF can limit the number of random steps (Northrup et al., 2013; Thurfjell et al., 2014). However, our VHF tracking methods provides a temportally coarse dataset, allowing us to generate a greater number of random steps enabling a broad sampling of the landscape and ensuring rare habitats were not missed. We therefore opted to generate 200 random steps for each observed step (i.e., relocation).

We created ten models including step length and turning angle. One of the ten models, our null model, only incorporated step length and turning angle to predict movement. We created six models that used a single habitat feature to predict selection, and three models that were multi-factor models which included a combination of the three different habitat features to predict habitat selection (Table 3).

We expanded our analysis to investigate habitat selection at the population level, including all of our telemetered snakes aside from PYBI060 who did not leave the forest and therefore did not utilize habitat features found within the agricultural matrix. We adapted code provided by Muff et al. (2019b) who used Poisson model with stratum‐specific effects to generate population level step-selection models. We created six single factor models using the same habitat features and rasters used for exploring individual habitat selection (i.e., inverted distance to forest, settlements, roads, water bodies, aquatic agriculture, and terrestrial agriculture), with individual random intercepts and slopes. As with individual models we randomly generated 200 steps for each observed step. We used a prior precision of 0.0001 for all fixed effects as used by Muff et al. (2019). These models provided us with the amount of individual variation that exists between the selection of our different habitat types as well as population level estimates of habitat selection. We fitted these Bayesian models via integrated nested Laplace approximations using the *INLA* package v.20.03.17 (Rue et al., 2020). With all analyses, we have provided all R code with exact model specification at DOI: 10.5281/zenodo.4026928.

## Results

We tracked seven adult Burmese pythons (six females, one male) over the course of our study period (2018-09-29 to 2020-07-22) (Table 2). We set out to track only females; however, due to a misidentification of one individual’s sex (human error during processing), we tracked one adult male. We tracked individuals for a mean of 327 ± SE 85 days (range = 41 – 662 days). During this time, implanted snakes were located on average 310 ± 80 times (range = 41 – 631 times) with mean time lag, or time between data points, of 25.43 ± 0.32 hours (range = 8.55 – 452.77 hours). Burmese pythons were found in a mean of 112 ± 33 unique locations (range = 23 – 234 unique locations) and were stationary for approximately 5 days (125 ± 11.6 hours; range= 24.1 – 2010 hours) at a time.

**Table 2.**
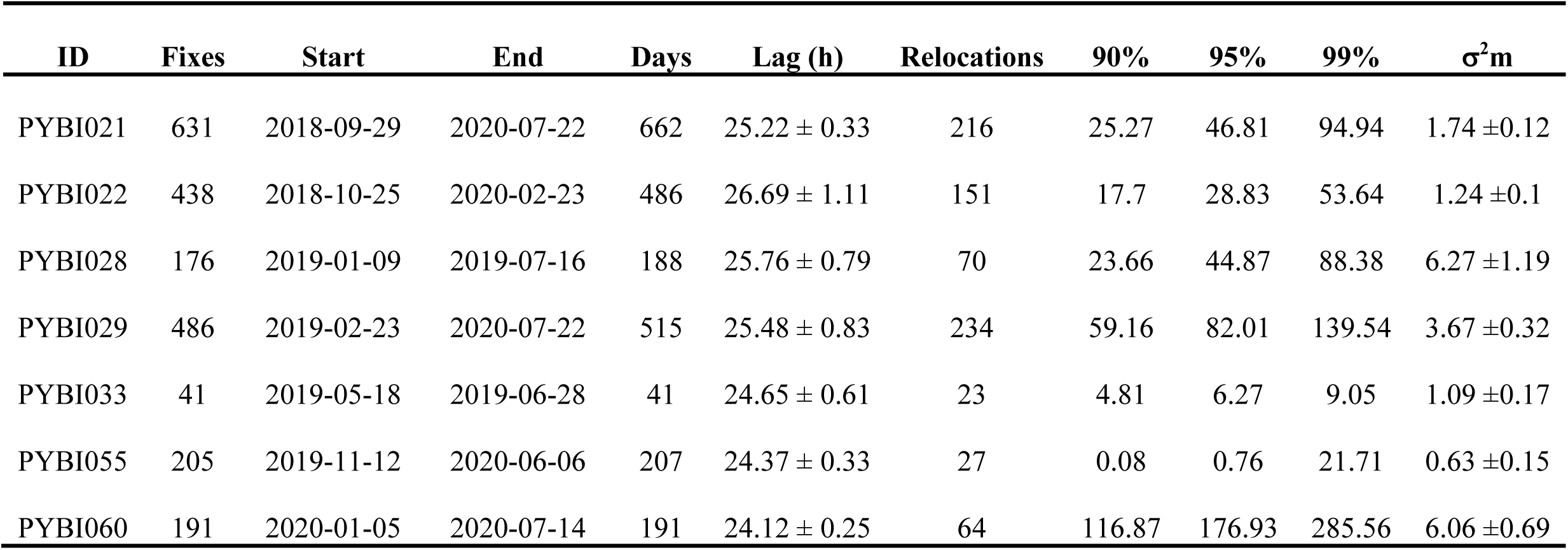
Tracking summary, 90%, 95% and 99% occurrence distributions (ha) and mean motion variance (σ^2^m) for radio tracked Burmese pythons.

### Occurrence distribution and motion variance

We calculated the occurrence distribution (i.e., estimate of uncertainty surrounding movement paths) for all tracked snakes (Table 2; Figure 3) using modified code from Marshall et al., (2020a; 2020b). On average, Burmese pythons 99% confidence area (99% contour generated from the dBBMM occurrence distribution) were 98.97 ± 35.42 ha. Excluding our single tracked male, female Burmese pythons had a mean 99% confidence area of 100.74 ± 41.86 ha. The largest 99% confidence area (285.56 ha) belonged to PYBI060 who was tracked exclusively within the protected core area of the biosphere reserve. The snake with the smallest 99% confidence area, PYBI033 (9.05 ha), was tracked for the shortest duration (41 days) due to an early transmitter failure. Our tracked male, tracked for over six months, had one of the smallest 99% confidence areas (88.38 ha). Our longest tracked snake, PYBI021 had a 99% confidence area of 99.94 ha. Movements within the 99% confidence areas showed site fidelity. Five of seven snakes returned to previously used shelter sites (range: 4-35 revisits); revisiting sites once every 43.47 ± 14.64 days on average.

**Figure 3.**
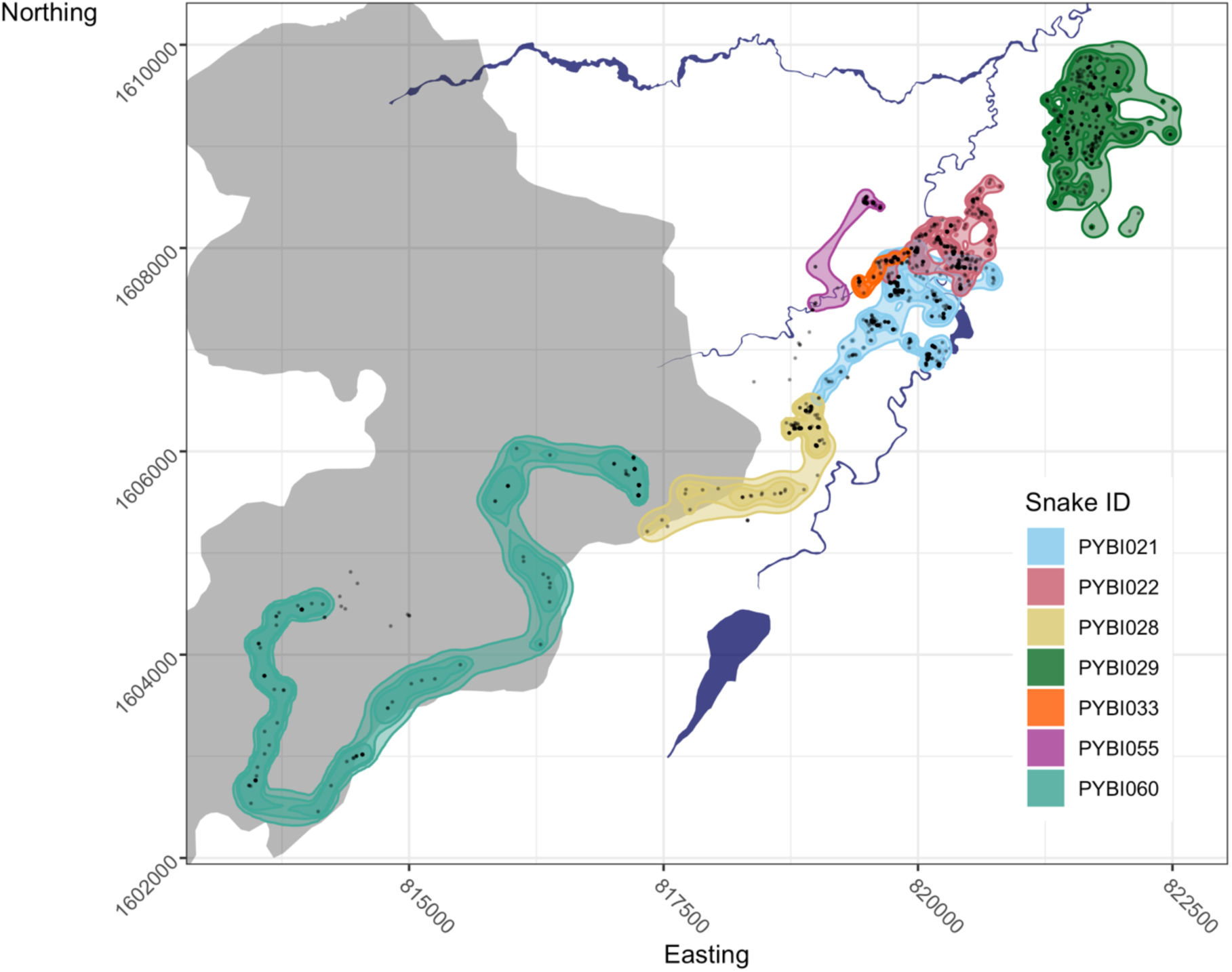
Dynamic Brownian Bridge Movement Models for all tracked snakes. Snake locations are shown by black points on the map. The grey shaded area marks the protected core area of the Sakaerat Biosphere Reserve. Major irrigation canals and reservoirs are shown by dark blue areas.

We also used dynamic Brownian and Bridge Movement Models to calculate the mean motion variance of our individuals (Figure 4). On average, our motion variance was low (mean = 2.66 ± SE 0.14 σ^2^m; range= 5.53^-05^ – 98.45 σ^2^m). The individuals that exhibited the highest mean motion variance were our tracked male, PYBI028 (6.27 ± 1.19 σ^2^m) and the female that remained entirely in the forest, PYBI060 (6.06 ± 0.69 σ^2^m). The lowest mean motion variance belonged to our largest tracked female, PYBI055, who remained stationary throughout much of her tracking duration during breeding and nesting (0.63 ± 0.15 σ^2^m).

**Figure 4.**
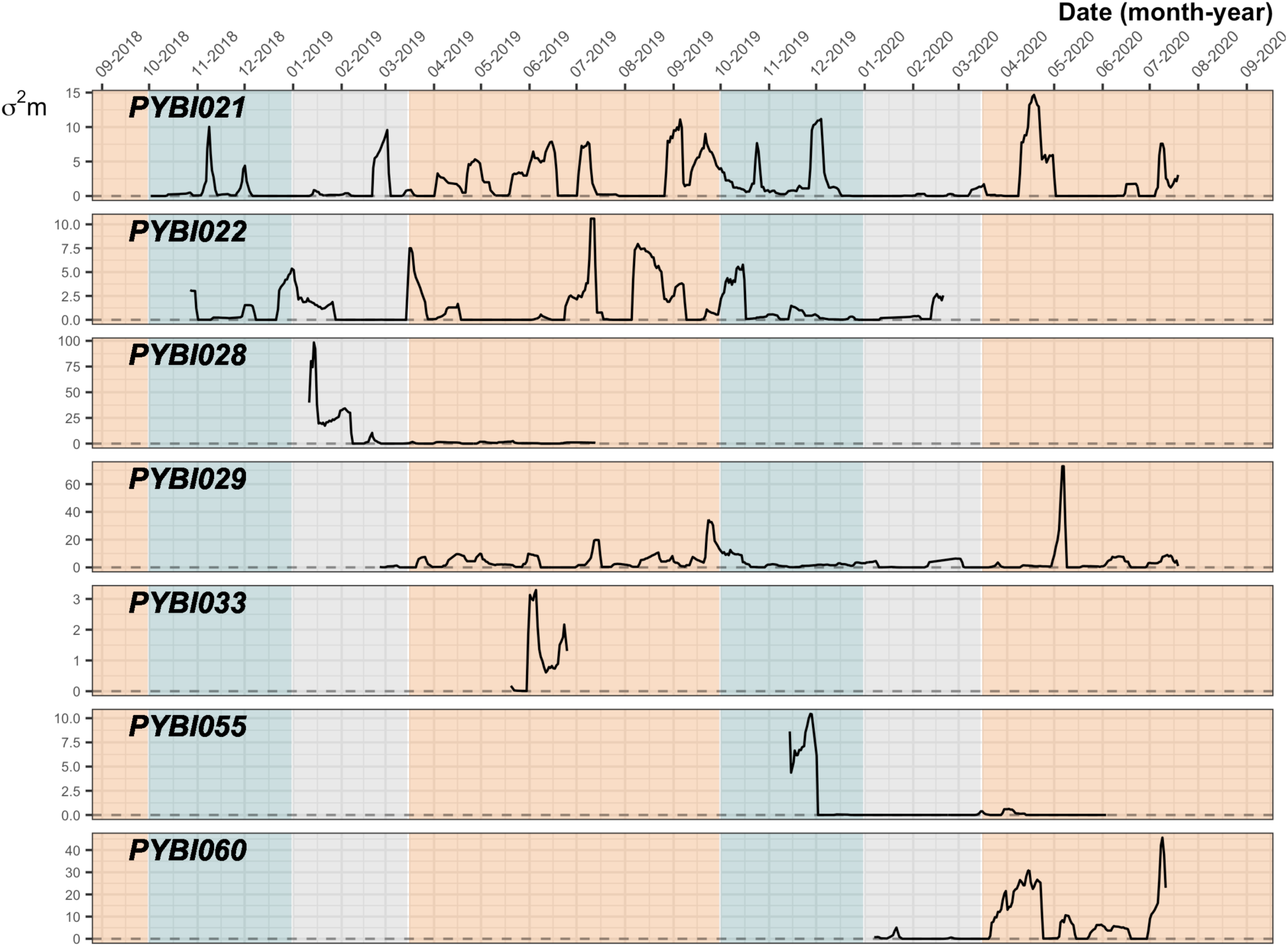
Motion variance for tracked females and one male (PYBI028) throughout individual tracking duration. Blue= wet season, grey = dry season, orange = hot season.

We were able to explore seasonal differences in motion variance for three individuals (PYBI021, PYBI022, and PYBI029) that we tracked for one year or longer. There appeared to be slight variation in movement depending on the season with mean motion variance highest during the hot season (3.12 ± 0.21 σ^2^m). Mean motion variance was slightly lower during the wet season (1.57 ± 0.12 σ^2^m) and lowest during the dry season (0.76 ± 0.09 σ^2^m). While we cannot fully explore seasonal differences in motion variance for our tracked male, his motion variance peaked in the dry season before greatly reducing his movement immediately prior to the hot season. In contrast, all tracked females tended to limit their movements, showing very few peaks in motion variance, during the dry season.

### Integrated step selection analysis

We created a total of ten models to explore the habitat selection of Burmese pythons in relation to certain habitat features such as forested areas, bodies of water and irrigation canals, human settlements, roads, and aquatic agriculture (Table 3; Figure 5). For our analysis, we excluded one individual, PYBI060, as she was our only snake that we tracked exclusively within the forested core area of the biosphere reserve (thereby having no opportunity to show attraction/avoidance of chosen landscape features). Habitat selection was best illustrated by four different models, Model 5, Model 6, Model 9 and Model 10. Of these models, Model 10 was the top model for three individuals (PYBI021, PYBI029 and PYBI033) and showed a positive association for the features included in the model. Model 10 was a multi-factor model that included distance to water, human settlements, and aquatic agriculture as predictors of movement. Model 6 was the top model for PYBI028 and was a single factor model showing movements as predicted by aquatic agriculture. Model 5 was a single factor model that investigated selection for distance to water bodies and was the top model for PYBI022. Model 9 was another multi-factor model that incorporated roads, terrestrial agriculture and water to predict selection and was the top model for PYBI055. For several of our models, our confidence intervals for particular individuals were very broad and in some cases overlapped zero, therefore limitting our inferences. We suspect that the uncertainity within our models is likely due to our coarse dataset and the fact that Burmese pythons relocated relatively infrequently.

**Table 3.** Model formulas and AIC scores for individual habitat selection models.

| Model | Model formula | PYBI021 | PYBI022 | PYBI028 | PYBI029 | PYBI033 | PYBI055 |
| --- | --- | --- | --- | --- | --- | --- | --- |
| model1 | log_sl*cos_ta+strata(step_id_) (null model) | 2238.4 | 1564.92 | 725.44 | 2456 | 228.63 | 269.39 |
| model2 | Model1 + forest + forest:sl + forest:ta | 2235.6 | 1569.43 | 721.25 | 2452.23 | 232.58 | 273.9 |
| model3 | Model1 + settle + settle:sl + settle:ta | 2233.67 | 1570.53 | 720.7 | 2458.89 | **\*228.09\*** | 269.77 |
| model4 | Model1 + road + road:sl + road:ta | 2234.94 | 1564.77 | 726.27 | 2456.14 | **\*227.85\*** | 267.63 |
| model5 | Model1 + water + water:sl + water:ta | 2221.18 | **\*1556.79\*** | 718.98 | **\*2438.1\*** | 228.75 | 258.01 |
| model6 | Model1 + aq.ag + aq.ag:sl + aq.ag:ta | 2236.23 | 1566.62 | **\*713.66\*** | 2460.39 | **\*227.18\*** | 268.8 |
| model7 | Model1 + terr.ag + terr.ag:sl + terr.ag:ta | 2237.94 | 1565.76 | 724.98 | 2453.68 | 230.82 | 272.42 |
| model8 | Model1+ road + forest + settle | 2240.91 | 1567.83 | 728.49 | 2444.86 | 232.65 | 271.58 |
| model9 | Model1+ road + terr.ag + water | 2219.99 | 1563.62 | 717.28 | **\*2439.77\*** | **\*227.46\*** | **\*252.34\*** |
| model10 | Model1 + water + settle + aq.ag | **\*2216.48\*** | 1562.47 | 723.76 | **\*2437.96\*** | **\*226.24\*** | 258.73 |

sl: step length, ta: turn angle, settle: human settlement, aq.ag: aquatic agriculture, terr.ag:
terrestrial agriculture. \* indicates AIC score within $< 2 \Delta$ AIC of model that best predicts
selection for individual.

**Figure 5.**
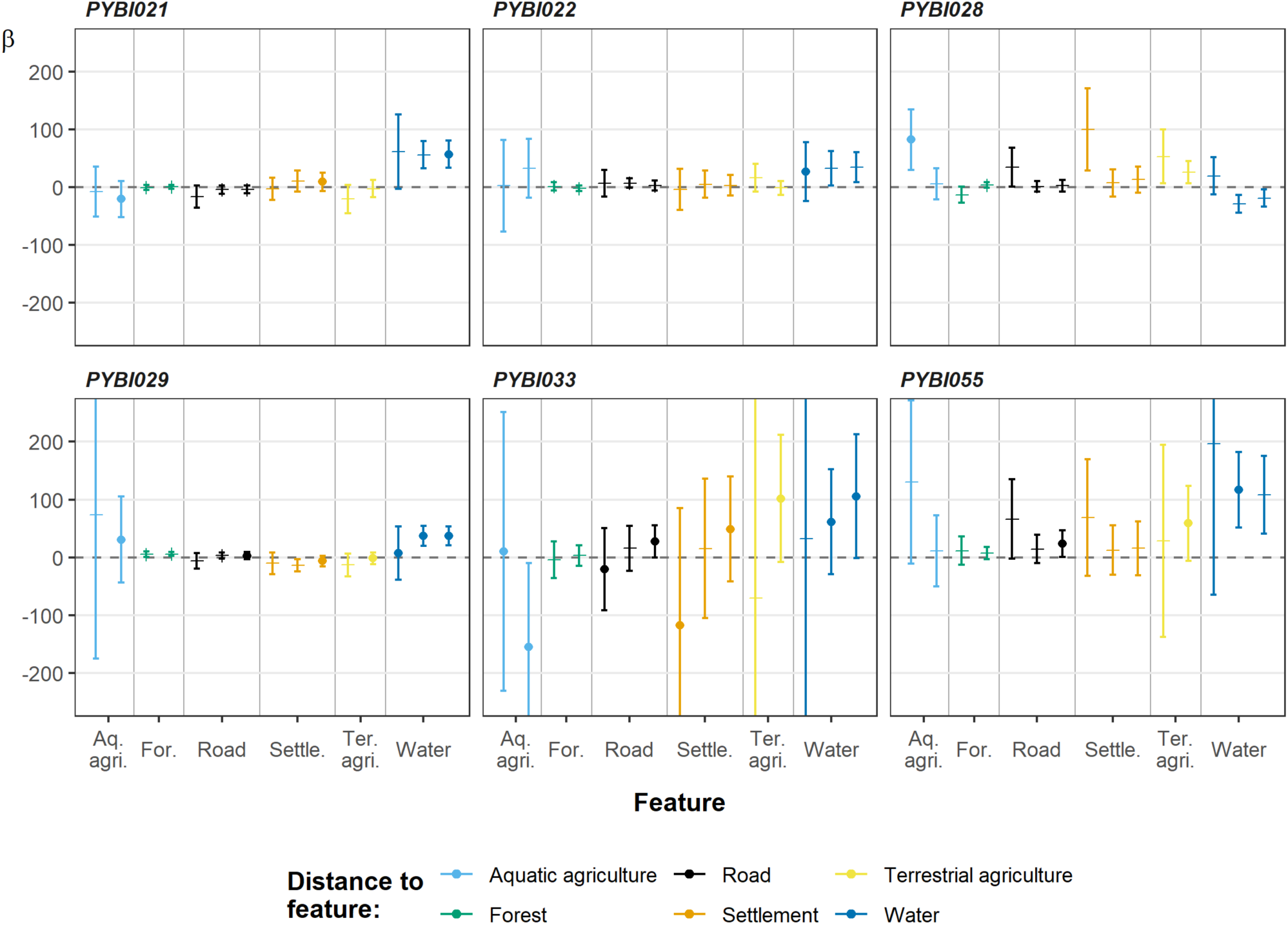
Habitat selection of individuals based on distance to habitat features. Positive estimates suggest association with habitat feature. Error bars indicate 95% confidence intervals. Circles mark the habitat features that were included in models with AIC scores within < 2 Δ AIC of top performing models.

Our estimates suggested that three out of six snakes were very minimally associated with forested areas. Instead, habitat features found within the agricultural matrix in our study site were better descriptors of habitat selection by Burmese pythons. For two of our snakes, PYBI028 and PYBI055, we saw a positive association with human settlements (*β* 99.94 95% CI [Confidence interval] 29.12 – 170.75, and *β* 68.89, 95% CI −31.78 – 169.57, respectively). Three snakes showed a slight association with terrestrial agriculture. Our tracked male, PYBI028, who was negatively associated with forested areas (*β* −13.13, 95% CI −26.77 – 0.49), showed a positive association with each habitat feature found within the agricultural landscape. All snakes were associated with either water bodies or aquatic agriculture. We did not detect consistent interaction between step length and distance to landscape features as there was pronounced individual heterogeneity (Supplementary Figure 2).

At the population level, mean estimates for habitat selection were low (range: −1.37^-04^ – 0.003) (Table 4; Figure 6). Reflecting findings from the individual step-selection models, the population models showed a positive association with water bodies (*β* 0.003, 95% CrI [Credible interval] 0.001 – 0.004). We failed to detect any active avoidance of human settlements or roads. At the population level, interaction between step length and distance to habitat feature was consistently low (Supplementary Figure 3). There appeared to be low variation between individuals for each model, which is to be expected with our sampling regime and coarse data set.

**Table 4.** Model formulas, Mean estimates (β) and the amount of variation between individuals for habitat selection at the population level.

| Model | Model formula | $\beta$ | Individual Variance |
| --- | --- | --- | --- |
| 1 | Step_id + forest + forest:sl + forest:ta | 8.04E-04 | 4.91E-06 |
| 2 | Step_id + settle + settle:sl + settle:ta | 5.66E-04 | 4.83E-06 |
| 3 | Step_id + road + road:sl + road:ta | 0.001419404 | 9.34E-06 |
| 4 | Step_id + water + water:sl + water:ta | 0.002720069 | 6.10E-12 |
| 5 | Step_id + aq.ag + aq.ag:sl + aq.ag:ta | -1.37E-04 | 1.18E-05 |
| 6 | Step_id + terr.ag + terr.ag:sl + terr.ag:ta | 0.004166886 | 8.17E-06 |

sl: step length, ta: turn angle, settle: human settlement, aq.ag: aquatic agriculture, terr.ag: terrestrial agriculture

**Figure 6.**
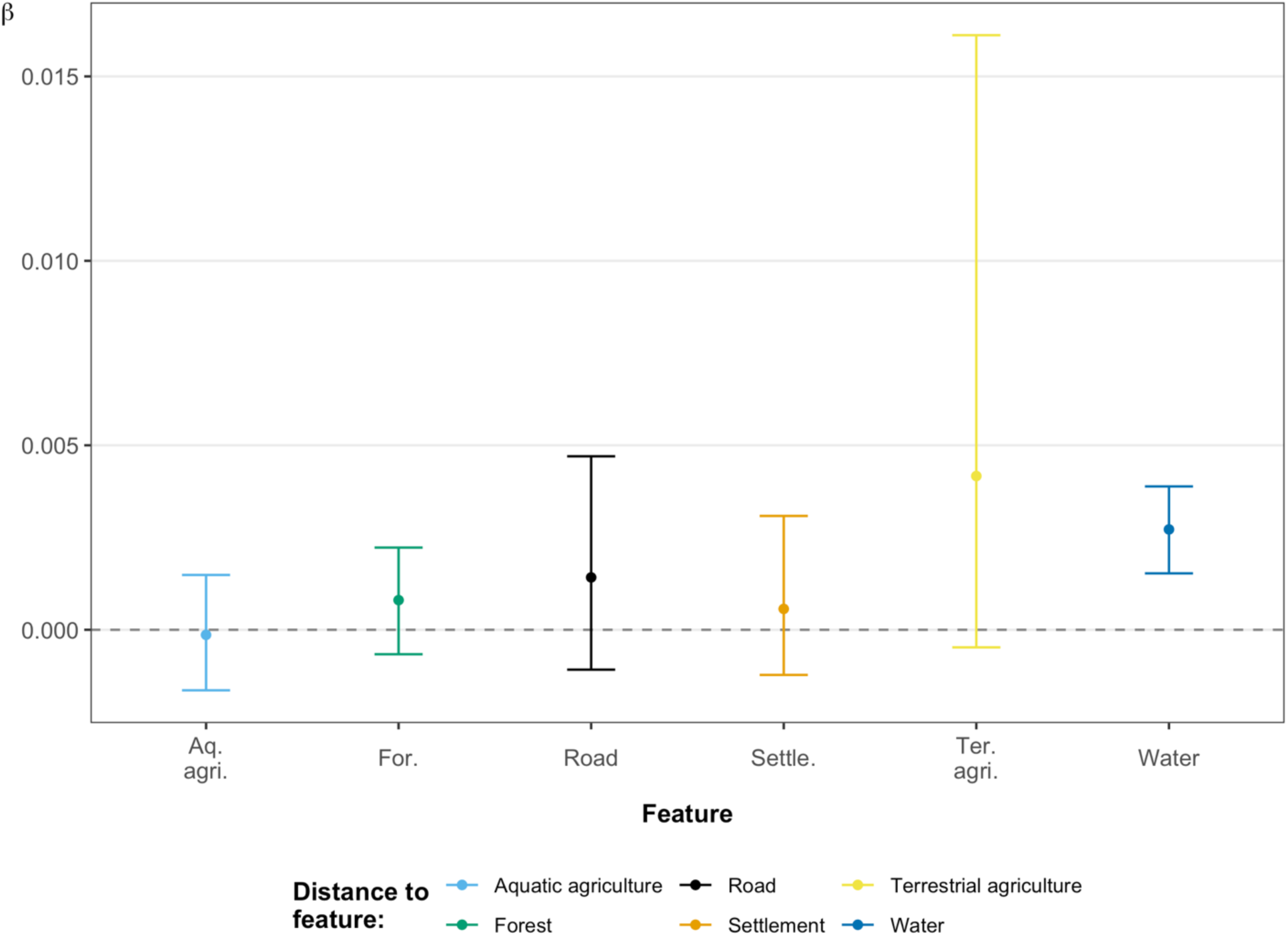
Habitat selection at the population level for all snakes tracked in the agricultural matrix. Selection is based on distance to habitat features with positive estimates indicating positive association. Error bars indicate 95% credible intervals.

## Discussion

Our study is one of the first to investigate the space use, movements, and habitat use of native Burmese pythons. Our results include the movements of several individuals that were concurrently tracked for extended durations (≥ 1 year) revealing seasonal movements. Burmese python movements covered small areas (mean= 98.97 ± 35.42 ha, 99% contour), with low motion variance (mean= 2.66 ± SE 0.14σ^2^m), and most exhibit site fidelity. Within our study site, all individuals (aside from one individual that never left the forest) showed movements associated with habitat features found within the agricultural matrix of the Biosphere reserve’s transitional zone. Of these habitat features, aquatic features (water-based agriculture such as rice paddies and water bodies) appeared to be selected for by all individuals to varying degrees.

Previous studies on Burmese pythons (both in their native range and invasive range) have used more “traditional” home range estimators, such as MCPs and KDEs, which are limited in their ability to be compared across studies (Silva et al., 2018; Silva et al., 2020). Rather, we measured the occurrence distributions of our tracked snakes during our tracking period. Despite this, we are able to draw some broad inferences between our results and those from studies in their invasive range. For instance, in their introduced range (Florida, USA), Burmese pythons appear to use very large areas and do not tend to show site fidelity, exhibiting large linear movements instead (Hart et al., 2015).

In contrast, the telemetered pythons in our study moved within much smaller areas. These differences in space use could be linked to the pressures that are present in our study site that may not be as prominent, or exist, in their invasive range. Our study site has been largely modified by agricultural conversion that can influence the movements and behaviors of animals (Doherty & Driscoll, 2018; Marshall et al., 2020; Tucker et al., 2018). Burmese pythons may alter their movements and behaviors in a human dominated landscape in attempt to avoid conflict with humans or other anthropogenic risks (Ditchkoff et al., 2006; Doherty et al., 2019; Wang et al., 2017). Furthermore, we have observed King Cobras preying upon Burmese pythons within the agricultural landscape of our study site. Animals may alter or reduce movement in attempt to avoid predation (Doherty et al., 2019; Rettie & Messier, 2001; Sih, 1984), which could partially explain low motion variance exhibited by our telemetered Burmese pythons.

Recursive analysis showed Burmese pythons frequently reuse shelter sites, suggesting that these sites may help to form what might be considered a “consistent” home range or occurrence distribution. Interestingly, the forest-dwelling individual from within the protected area of the biosphere reserve never reused a shelter site. The frequency of site reuse from individuals tracked within the agricultural matrix suggests that these snakes may more carefully select for and subsequently reuse shelter sites in areas highly modified by human activity. In areas or times of year with few sufficient shelter sites animals may repeatedly return to suitable sites to take refuge (Beck & Jennings, 2003; Whitaker & Shine, 2003; Young et al., 2017). The only other individual tracked within the forest was our male (PYBI028) who briefly moved into a patch of forest during what we assume to be the breeding season (December-March) for Burmese pythons at our study site.

Consistently, our snakes exhibited low mean motion variances and typically showed intermittent activity peaks followed by extended periods of the snake remaining stationary. Burmese pythons can be characterized as ambush predators (Ross & Winterhalder, 2015), so these peaks in activity we see could be linked to our snakes searching for new ambush sites and then remaining there to capture and digest a meal. Aside from snakes that were captured after consuming livestock, no observations of snakes with evident food items (visible bulge caused by ingested prey) were made during the study. Although infrequent, we observed molted skin outside of snake shelter sites, suggesting that ecdysis may additionally explain prolonged stationary periods.

For reproductive females, long durations of inactivity can also be linked to reproduction as Burmese pythons are known to invest a high level of maternal care and will remain coiled around their eggs following oviposition until hatching (Krysko et al., 2008; Snow et al., 2010). Two of our snakes (PYBI022 and PYBI055), nested between April and June and dramatically limited movement during brooding. Interestingly, PYBI055 remained in the same burrow for 84 days between late December through early to mid-March prior to moving to her oviposition site. We observed several conspecifics entering and leaving the burrow throughout this 84-day period, which we assume were males. Burmese pythons are known to form mating aggregations in their invasive range (Smith et al., 2016); which is what we suspect was also occurring within this burrow. When we examine the movements of PYBI022 during this similar timeframe, we see that there were several locations she visited where she would remain for an extended period of time (>14 days) but did not exceed 21 days in a single location.

Universally, Burmese pythons showed a positive association with aquatic habitat features, whether that was aquatic agriculture (i.e., rice paddy) or water bodies (i.e., ponds, irrigation canals). In contrast, three of our snakes showed a slight attraction to terrestrial agriculture. Terrestrial agriculture, such as cassava and orchards, may not be selected for because it lacks suitable shelter or microclimate for ectotherms such as the Burmese python (Frishkoff et al., 2015; Gallmetzer & Schulze, 2015). In their invasive range, Burmese pythons regularly use aquatic environments and are thought to be semi-aquatic (Hunter et al., 2015; Mazzotti et al., 2011), which is consistent with our findings of Burmese pythons selecting for aquatic features.

We are unable to identify the underlying drivers behind Burmese python’s selection of aquatic features; however, one explanation could be the abundance of potential prey items in aquatic environments, such as wading birds which we frequently observed during radio-tracking. Rice paddy fields often act as wetland-like habitat and support communities of wading birds (Fujioka et al., 2010; Lawler, 2001). The initial capture of our first tracked individual (PYBI021) occurred after a group of forestry department workers witnessed her consume several domestic ducks in a small man-made pond. Aquatic environments and the vegetation that typically grows on the edges of ponds and irrigation canals could also serve as suitable refuge in an area highly modified by human settlements, roads, and agricultural practices. Furthermore, irrigation canals could also serve as movement corridors for Burmese pythons as seen with King Cobras that often travel along irrigational canals while in the agricultural land within our study site (Marshall 2020b; Marshall et al., 2019). The use of connected systems of aquatic agriculture in their native range (this study site, rice paddies and irrigation canals, Thailand) may help explain their success in their invasive range (natural everglade wetlands, Florida, USA) that is similarly characterized by interconnected wetlands.

Of our snakes that we tracked in the agricultural areas, only one strongly avoided human settlements, which is concerning from a human-wildlife conflict perspective. Human wildlife conflict is likely to occur when wildlife and humans compete for resources, space, and when wildlife causes economic loss through damages or livestock loss (Barua et al., 2013; Dickman, 2010). Within our study site, many residents keep livestock such as chickens, ducks, and geese. On two occasions, we initially captured study animals after they were seen consuming livestock animals. We captured a third Burmese python after she became entangled in netting material used to enclose a chicken coop; however, she was unsuccessful in her apparent attempt to consume livestock. The nature of these captures serves as evidence that Burmese pythons do initiate human-wildlife conflict via consumption of livestock in our study site. We were fortunate that none of these instances led to the persecution of Burmese pythons, as direct human-snake conflict results in snake mortality for other species in our study site (Crane et al., 2016; Marshall et al., 2018). It is difficult to determine whether Burmese pythons fall victim to persecution in our study site as we suspect that people would hesitate to notify us about killing or harming a protected species.

While we observed patterns of movement, space use and habitat selection that were fairly uniform across our sample, we acknowledge that our inferences are limited due to several biases that may affect our study animals, highlighted by Webster and Rutz (2020). These biases include: Acclimation & habituation, Trappability & self-selection and Genetic make-up.

Acclimation & habituation: We attempted to minimize disturbance to Burmese pythons by limiting tracks to one time per day and by tracking them during periods of inactivity (i.e., during the day). However, our tracking protocol included directly homing in on snakes, it is possible that approaching the snakes could have altered their behaviors and movements over time.

Trappability and self-selection: Our sampling method was highly dependent on notification of Burmese pythons from local residents, meaning that our sample likely included animals that were more likely to enter human settlements. We stress that this sample is non-random and thus likely not representative of the overall population. We strongly caution extrapulating outside of the sample. We saw variation in movements and space use between our animals tracked in human dominated areas compared to our one individual that never left the forest, suggesting that our results may not be generalizable to Burmese pythons in different populations.

Genetic make-up: We do not currently have information about the genetic make-up and diversity within our sample size. We tracked several snakes that were captured within close proximity to eachother and could have been related. It is possible that there is little genetic variation in our sample, meaning that our results may not be applicable to more genetically diverse populations of Burmese pythons.

Our inferences are limited by the biases mentioned above. Despite sample limitations we revealed that non forest pythons showed general selection for aquatic systems and water bodies, supporting hypotheses for Burmese pythons success in their invasive range (Mutascio et al., 2018). Due to our snakes perceived indifference to human dominated spaces, we suggest that future research target the interactions between humans and pythons within an anthropogenically altered landscape. Furthermore, we suggest expanding studies to explore male movements, and how Burmese pythons move in forested or natural areas; studies of this type would help gauge the generalizablity of our findings and how predicted habitat loss may impact Burmese pythons.

## Conclusion

Our study serves as one of the first to investigate the ecology of native Burmese pythons and has provided us with estimates of their space use, movements and habitat selection in a landscape heavily modified by human activities. In agricultural land, Burmese pythons tend to move across small areas and appear to limit their movement by spending long periods of time stationary and making small, infrequent moves often associated with water. We suspect that this limited movement influences detection probablity of Burmese pythons (Steen, 2010). Burmese pythons tracked in agricultural areas returned to previously used shelter sites on several occasions, which we presume is linked to the necessity for careully selecting shelter in areas heavily modified by anthropogenic practices.

## Research permission

Our research was permitted by National Research Council of Thailand (NRCT) and by the National Park, Wildlife and Plant Conservation Department, Thailand (DNP) (0002/6115). Our methodology is in accordance with the Ethical Principles and Guidelines for the Use of Animals for Scientific Purposes provided by the National Research Council of Thailand. All work was conducted under Institute of Animals for Scientific Purpose Development (IAD) licensing belonging to C.T.S. We were approved to study Burmese pythons in the Sakaerat Biosphere Reserve by Thailand Institute of Scientific and Technological Research and the Sakaerat Environmental Research Station.

## Acknowledgments

We thank Nakhon Ratchasima Zoo for facilitating radio-transmitter implantation surgeries under the supervision and expertise of their veterinarians. We thank both the National Research Council of Thailand and the National Park, Wildlife and Plant Conservation Department, Thailand for permitting our research on Burmese Pythons. We thank Suranaree University of Technology, School of Biology for supervising our research and assisting with logistics. We thank Pluemjit Boonpueng for providing assistance with general logistics and paperwork. We thank the Institute of Animals for Scientific Purpose Development for providing animal use licenses for C.T.S. We are grateful for previous project leaders from the Sakaerat Conservation and Snake Education Team lending us equipment to conduct our research. We thank the Thailand Institute of Scientific Research for providing partial funding as well as logistical support. We thank the Sakaerat Environmental Research Station for supporting us, housing us, and assisting with logistics. We thank Nithina Kaewtongkum and Kanoktip Somsiri for facilitating snake calls. We enthusiastically thank the residents and community of Udom Sab for allowing us to track study animals throughout their land. We commend the Udom Sab/Hook 31 Rescue team for working towards human-snake conflict mediation and thank them for assisting in the capture of several Burmese pythons. We are grateful for Joli Stavish, Shannon Thrasher and several members of the King Cobra Telemetry Project for their hard work, enthusiasm and sense of humor during the tracking of Burmese pythons throughout our study.

## Author contributions

*Conceptualization:* S.N.S., C.T.S., B.M.M., G.A.G., and M.D.J; *Methodology*: S.N.S., C.T.S., B.M.M., G.A.G., and M.D.J; *Formal Analysis:* S.N.S., B.M.M., *Investigation:* S.N.S., M.D.J.; *Resources*: S.W.; *Supervision:* S.W., C.T.S., G.A.G.; *Visualization*: S.N.S., M.D.J., B.M.M.; *Writing – Original Draft Preparation:* S.N.S.; *Writing – Review & Editing:* S.N.S., M.D.J., B.M.M., C.T.S., G.A.G., S.W.

