## Supplementary figures and tables for "Restricted constrictors: Space use and habitat selection of native Burmese pythons in Northeast Thailand"

**Supplementary Table 1.** Capture and release dates, coordinates, and methods for all radio-tracked snakes.

| Snake ID | Capture date | Capture (E) | Capture (N) | Capture Method | Release date | Release (E) | Release (N) | Distance (m) |
|---|---|---|---|---|---|---|---|---|
| PYBI021 | 2018-09-24 | 819617 | 1607440 | Notation | 2018-09-28 | 819808 | 1607405 | 193 |
| PYBI022 | 2018-10-18 | 820364 | 1608058 | Notation | 2018-10-24 | 820375 | 1608002 | 57 |
| PYBI028 | 2019-01-04 | 819044 | 1606437 | Opportunistic | 2019-01-08 | 819308 | 1606727 | 393 |
| PYBI029 | 2019-01-31 | 821671 | 1609646 | Notation | 2019-02-22 | 821671 | 1609646 | 0 |
| PYBI033 | 2019-05-10 | 819471 | 1607766 | Notation | 2019-05-17 | 819418 | 1607720 | 70 |
| PYBI055 | 2019-11-05 | 818462 | 1606647 | Notation | 2019-11-12 | 818477 | 1606766 | 121 |
| PYBI060 | 2019-12-26 | 817051 | 1605783 | Opportunistic | 2020-01-05 | 817241 | 1605873 | 210 |

(E) and (N) refer to UTMs Easting and Northing.

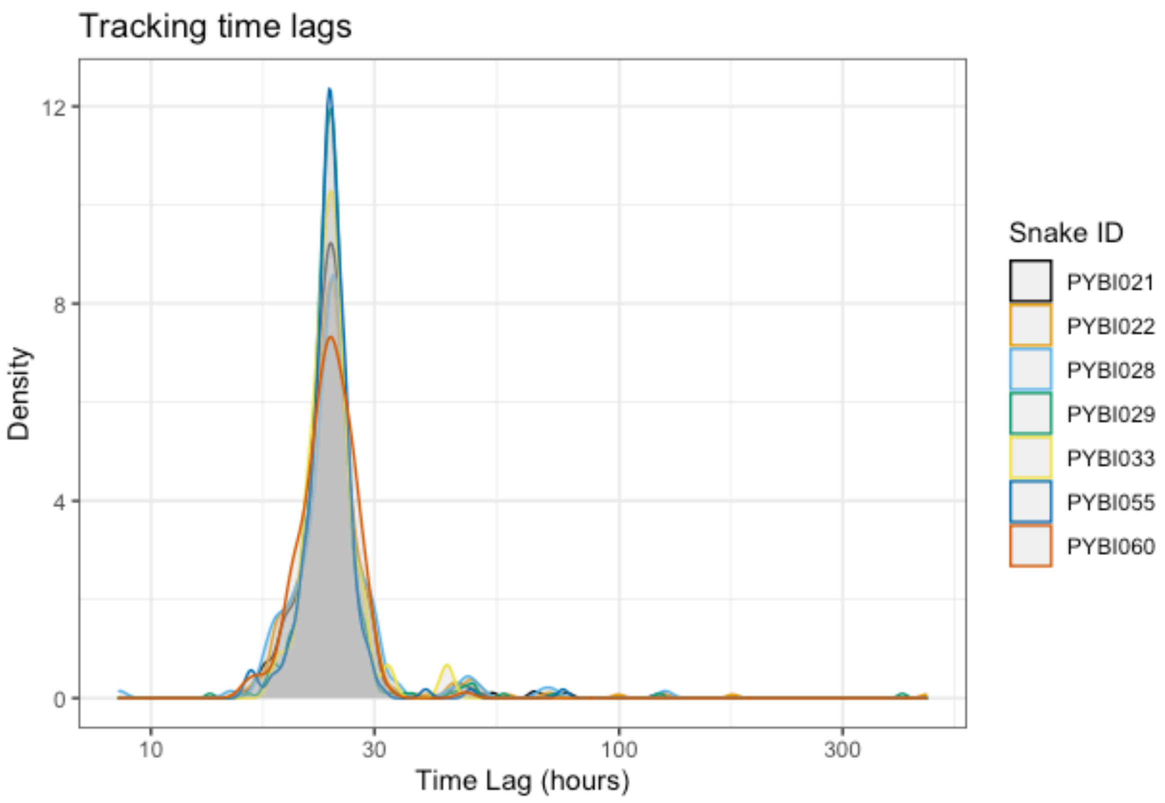


**Supplementary Figure 1.** Density of time lag between tracks for all individuals.

**Supplementary Table 2.** All integrated step selection analysis results from individual habitat selection.

| term | estimate | std.error | statistic | p.value | conf.low | conf.high | id | model | AIC |
|---|---|---|---|---|---|---|---|---|---|
| log_sl | -0.0202182 | 0.05279295 | -0.3829724 | 0.70174024 | -0.1236905 | 0.08325405 | PYBI021 | model1 | 2238.39539 |
| cos_ta | -0.7006985 | 0.30182713 | -2.3215224 | 0.02025866 | -1.2922688 | -0.1091282 | PYBI021 | model1 | 2238.39539 |
| log_sl:cos_ta | 0.17594274 | 0.07312053 | 2.4062017 | 0.01611936 | 0.03262914 | 0.31925634 | PYBI021 | model1 | 2238.39539 |
| dist_forest | -0.7797137 | 2.19599122 | -0.3550623 | 0.72254291 | -5.0837774 | 3.52434999 | PYBI021 | model2 | 2235.60231 |
| log_sl | -3.942199 | 2.31074058 | -1.7060327 | 0.08800199 | -8.4711673 | 0.58676931 | PYBI021 | model2 | 2235.60231 |
| cos_ta | 11.4486933 | 4.94707995 | 2.31423252 | 0.02065497 | 1.75259474 | 21.1447918 | PYBI021 | model2 | 2235.60231 |
| dist_forest:log_sl | 0.52047042 | 0.30687779 | 1.69601854 | 0.08988237 | -0.080999 | 1.12193984 | PYBI021 | model2 | 2235.60231 |
| dist_forest:cos_ta | -1.6190168 | 0.65812389 | -2.4600487 | 0.01389182 | -2.9089159 | -0.3291177 | PYBI021 | model2 | 2235.60231 |
| log_sl:cos_ta | 0.19473642 | 0.07331735 | 2.6560754 | 0.00790559 | 0.05103705 | 0.3384358 | PYBI021 | model2 | 2235.60231 |
| dist_settle | -3.2158744 | 9.75831317 | -0.3295523 | 0.74173828 | -22.341817 | 15.9100679 | PYBI021 | model3 | 2233.6715 |
| log_sl | -33.483572 | 13.7288481 | -2.4389207 | 0.0147312 | -60.39162 | -6.5755243 | PYBI021 | model3 | 2233.6715 |
| cos_ta | 59.8001653 | 29.4211492 | 2.03255709 | 0.04209729 | 2.13577262 | 117.464558 | PYBI021 | model3 | 2233.6715 |
| dist_settle:log_sl | 3.77920169 | 1.55081432 | 2.43691435 | 0.01481319 | 0.73966146 | 6.81874191 | PYBI021 | model3 | 2233.6715 |

| | | | | | | | | | |
|---|---|---|---|---|---|---|---|---|---|
| dist_settle:cos_ta | -6.8378614 | 3.32497034 | -2.056518 | 0.03973261 | -13.354683 | -0.3210393 | PYBI021 | model3 | 2233.6715 |
| log_sl:cos_ta | 0.19679221 | 0.07321787 | 2.68776181 | 0.00719327 | 0.05328781 | 0.3402966 | PYBI021 | model3 | 2233.6715 |
| dist_road | -16.122819 | 9.91713072 | -1.6257544 | 0.10400192 | -35.560038 | 3.31440041 | PYBI021 | model4 | 2234.93798 |
| log_sl | -27.046939 | 15.6996261 | -1.722776 | 0.08492904 | -57.817641 | 3.72376244 | PYBI021 | model4 | 2234.93798 |
| cos_ta | 84.8967001 | 31.9839535 | 2.65435291 | 0.00794607 | 22.2093031 | 147.584097 | PYBI021 | model4 | 2234.93798 |
| dist_road:log_sl | 3.44970083 | 2.00415835 | 1.72127159 | 0.08520155 | -0.4783774 | 7.37777902 | PYBI021 | model4 | 2234.93798 |
| dist_road:cos_ta | -10.933208 | 4.08510677 | -2.676358 | 0.00744271 | -18.93987 | -2.9265461 | PYBI021 | model4 | 2234.93798 |
| log_sl:cos_ta | 0.19551574 | 0.07381046 | 2.64888928 | 0.00807568 | 0.0508499 | 0.34018159 | PYBI021 | model4 | 2234.93798 |
| dist_water | 61.3838945 | 32.9641944 | 1.86213847 | 0.06258357 | -3.2247392 | 125.992528 | PYBI021 | model5 | 2221.17615 |
| log_sl | 7.72064314 | 60.7928908 | 0.12699911 | 0.8989411 | -111.43123 | 126.87252 | PYBI021 | model5 | 2221.17615 |
| cos_ta | 61.6967995 | 114.505698 | 0.53880986 | 0.59001806 | -162.73024 | 286.123844 | PYBI021 | model5 | 2221.17615 |
| dist_water:log_sl | -0.8738034 | 6.89134771 | -0.1267972 | 0.89910094 | -14.380597 | 12.63299 | PYBI021 | model5 | 2221.17615 |
| dist_water:cos_ta | -7.077632 | 12.9781907 | -0.5453481 | 0.58551411 | -32.514418 | 18.3591544 | PYBI021 | model5 | 2221.17615 |
| log_sl:cos_ta | 0.19773297 | 0.0748163 | 2.64291309 | 0.00821961 | 0.05109572 | 0.34437021 | PYBI021 | model5 | 2221.17615 |
| dist_aq.ag | -7.5683897 | 22.0253005 | -0.3436225 | 0.73113017 | -50.737185 | 35.600406 | PYBI021 | model6 | 2236.23475 |
| log_sl | 38.2602767 | 33.4522051 | 1.14372959 | 0.2527358 | -27.30484 | 103.825394 | PYBI021 | model6 | 2236.23475 |
| cos_ta | -137.62986 | 65.0898352 | -2.1144601 | 0.03447599 | -265.20359 | -10.056129 | PYBI021 | model6 | 2236.23475 |
| dist_aq.ag:log_sl | -4.0060551 | 3.50101838 | -1.1442542 | 0.25251822 | -10.867925 | 2.85581486 | PYBI021 | model6 | 2236.23475 |
| dist_aq.ag:cos_ta | 14.3286058 | 6.81148304 | 2.10359561 | 0.03541374 | 0.97834437 | 27.6788672 | PYBI021 | model6 | 2236.23475 |
| log_sl:cos_ta | 0.18681099 | 0.07377316 | 2.53223517 | 0.0113338 | 0.04221825 | 0.33140373 | PYBI021 | model6 | 2236.23475 |
| dist_terr.ag | -20.646123 | 12.4551158 | -1.657642 | 0.09738974 | -45.057702 | 3.76545511 | PYBI021 | model7 | 2237.9397 |
| log_sl | -37.329041 | 17.4790906 | -2.1356398 | 0.03270879 | -71.587429 | -3.0706527 | PYBI021 | model7 | 2237.9397 |
| cos_ta | 82.2276783 | 46.2088628 | 1.77947851 | 0.07516134 | -8.3400285 | 172.795385 | PYBI021 | model7 | 2237.9397 |
| dist_terr.ag:log_sl | 4.52776133 | 2.12176804 | 2.1339568 | 0.03284632 | 0.36917239 | 8.68635028 | PYBI021 | model7 | 2237.9397 |
| dist_terr.ag:cos_ta | -10.06951 | 5.61172019 | -1.7943714 | 0.07275391 | -21.06828 | 0.92925934 | PYBI021 | model7 | 2237.9397 |
| log_sl:cos_ta | 0.19250606 | 0.07300832 | 2.63676872 | 0.00836999 | 0.04941238 | 0.33559974 | PYBI021 | model7 | 2237.9397 |
| dist_road | -4.2306909 | 3.63022512 | -1.1654073 | 0.24385416 | -11.345801 | 2.88441956 | PYBI021 | model8 | 2240.91342 |
| dist_forest | 0.51781056 | 1.83208439 | 0.28263466 | 0.7774569 | -3.0730089 | 4.10862998 | PYBI021 | model8 | 2240.91342 |
| dist_settle | 10.6684866 | 9.33427714 | 1.14293656 | 0.25306493 | -7.6263604 | 28.9633336 | PYBI021 | model8 | 2240.91342 |
| log_sl | -0.0138324 | 0.05302224 | -0.2608792 | 0.79418566 | -0.1177541 | 0.09008929 | PYBI021 | model8 | 2240.91342 |
| cos_ta | -0.7149013 | 0.30298637 | -2.3595164 | 0.01829877 | -1.3087437 | -0.1210589 | PYBI021 | model8 | 2240.91342 |
| log_sl:cos_ta | 0.18163336 | 0.0734754 | 2.47202946 | 0.01343484 | 0.03762422 | 0.3256425 | PYBI021 | model8 | 2240.91342 |
| dist_road | -4.1608029 | 3.49857236 | -1.1892859 | 0.23432717 | -11.017879 | 2.69627292 | PYBI021 | model9 | 2219.98671 |
| dist_terr.ag | -2.4344319 | 7.68392676 | -0.3168213 | 0.75137918 | -17.494652 | 12.6257879 | PYBI021 | model9 | 2219.98671 |
| dist_water | 56.0128557 | 12.0371327 | 4.65333873 | 3.27E-06 | 32.4205092 | 79.6052023 | PYBI021 | model9 | 2219.98671 |
| log_sl | 0.01453099 | 0.05412152 | 0.26848815 | 0.7883236 | -0.0915452 | 0.12060722 | PYBI021 | model9 | 2219.98671 |
| cos_ta | -0.7535136 | 0.3063372 | -2.4597522 | 0.0139033 | -1.3539235 | -0.1531037 | PYBI021 | model9 | 2219.98671 |
| log_sl:cos_ta | 0.20187797 | 0.07459971 | 2.70614952 | 0.00680684 | 0.05566523 | 0.34809072 | PYBI021 | model9 | 2219.98671 |
| dist_water | 56.9867599 | 12.0573055 | 4.7263263 | 2.29E-06 | 33.3548755 | 80.6186444 | PYBI021 | model10 | 2216.47897 |
| dist_settle | 9.26322559 | 8.23683455 | 1.12460989 | 0.26075438 | -6.8806735 | 25.4071247 | PYBI021 | model10 | 2216.47897 |

| | | | | | | | | | |
|---|---|---|---|---|---|---|---|---|---|
| dist_aq.ag | -20.242022 | 15.9346909 | -1.2703116 | 0.20397367 | -51.473443 | 10.9893982 | PYBI021 | model10 | 2216.47897 |
| log_sl | 0.02402524 | 0.05464249 | 0.43968047 | 0.66016855 | -0.0830721 | 0.13112255 | PYBI021 | model10 | 2216.47897 |
| cos_ta | -0.7764358 | 0.30858792 | -2.5160925 | 0.0118664 | -1.381257 | -0.1716146 | PYBI021 | model10 | 2216.47897 |
| log_sl:cos_ta | 0.21009551 | 0.07528521 | 2.79066104 | 0.00526005 | 0.0625392 | 0.35765182 | PYBI021 | model10 | 2216.47897 |
| log_sl | 0.00687405 | 0.07502795 | 0.09161982 | 0.92700011 | -0.140178 | 0.15392613 | PYBI022 | model1 | 1564.92228 |
| cos_ta | -0.0561286 | 0.43750585 | -0.1282923 | 0.89791769 | -0.9136243 | 0.80136711 | PYBI022 | model1 | 1564.92228 |
| log_sl:cos_ta | -6.96E-04 | 0.10588688 | -0.0065712 | 0.99475698 | -0.2082303 | 0.20683867 | PYBI022 | model1 | 1564.92228 |
| dist_forest | 1.38935065 | 3.55881777 | 0.39039668 | 0.69624324 | -5.585804 | 8.3645053 | PYBI022 | model2 | 1569.43138 |
| log_sl | 2.16156858 | 4.3808922 | 0.4934083 | 0.62172412 | -6.4248224 | 10.7479595 | PYBI022 | model2 | 1569.43138 |
| cos_ta | 7.69971958 | 7.12657847 | 1.08042304 | 0.27995384 | -6.2681176 | 21.6675567 | PYBI022 | model2 | 1569.43138 |
| dist_forest:log_sl | -0.2898244 | 0.5887786 | -0.4922467 | 0.62254492 | -1.4438092 | 0.8641605 | PYBI022 | model2 | 1569.43138 |
| dist_forest:cos_ta | -1.0403936 | 0.95435384 | -1.090155 | 0.27564488 | -2.9108928 | 0.83010555 | PYBI022 | model2 | 1569.43138 |
| log_sl:cos_ta | -0.0058205 | 0.10605791 | -0.0548808 | 0.95623342 | -0.2136902 | 0.20204914 | PYBI022 | model2 | 1569.43138 |
| dist_settle | -3.7252969 | 18.2289486 | -0.2043616 | 0.83807094 | -39.45338 | 32.0027858 | PYBI022 | model3 | 1570.53409 |
| log_sl | -8.7616823 | 28.9445613 | -0.3027057 | 0.76211419 | -65.49198 | 47.9686153 | PYBI022 | model3 | 1570.53409 |
| cos_ta | -24.379728 | 47.9957247 | -0.5079562 | 0.61148403 | -118.44962 | 69.6901639 | PYBI022 | model3 | 1570.53409 |
| dist_settle:log_sl | 0.98989273 | 3.26758701 | 0.30294304 | 0.76193327 | -5.4144601 | 7.39424559 | PYBI022 | model3 | 1570.53409 |
| dist_settle:cos_ta | 2.74587604 | 5.41815885 | 0.50679135 | 0.61230123 | -7.8735202 | 13.3652723 | PYBI022 | model3 | 1570.53409 |
| log_sl:cos_ta | -0.0011952 | 0.1060606 | -0.0112687 | 0.99100911 | -0.2090701 | 0.2066798 | PYBI022 | model3 | 1570.53409 |
| dist_road | 6.65780479 | 11.9418039 | 0.55752086 | 0.57717161 | -16.747701 | 30.0633103 | PYBI022 | model4 | 1564.77401 |
| log_sl | -1.3960659 | 19.3090826 | -0.072301 | 0.94236237 | -39.241172 | 36.4490406 | PYBI022 | model4 | 1564.77401 |
| cos_ta | 62.595119 | 32.4253602 | 1.93043712 | 0.0535527 | -0.9574191 | 126.147657 | PYBI022 | model4 | 1564.77401 |
| dist_road:log_sl | 0.18033553 | 2.46380482 | 0.07319392 | 0.94165181 | -4.6486332 | 5.00930425 | PYBI022 | model4 | 1564.77401 |
| dist_road:cos_ta | -7.9902295 | 4.13446361 | -1.9325916 | 0.05328653 | -16.093629 | 0.11317024 | PYBI022 | model4 | 1564.77401 |
| log_sl:cos_ta | -0.0081109 | 0.10748127 | -0.0754631 | 0.93984628 | -0.2187703 | 0.20254856 | PYBI022 | model4 | 1564.77401 |
| dist_water | 27.1743948 | 25.9266723 | 1.04812505 | 0.29458099 | -23.640949 | 77.9897387 | PYBI022 | model5 | 1556.78833 |
| log_sl | -31.591483 | 47.4978796 | -0.6651135 | 0.50597788 | -124.68562 | 61.5026507 | PYBI022 | model5 | 1556.78833 |
| cos_ta | 229.373599 | 93.4314246 | 2.45499413 | 0.01408869 | 46.2513715 | 412.495826 | PYBI022 | model5 | 1556.78833 |
| dist_water:log_sl | 3.58497846 | 5.38599501 | 0.66561117 | 0.50565966 | -6.9713778 | 14.1413347 | PYBI022 | model5 | 1556.78833 |
| dist_water:cos_ta | -26.016567 | 10.591908 | -2.4562682 | 0.01403883 | -46.776325 | -5.2568084 | PYBI022 | model5 | 1556.78833 |
| log_sl:cos_ta | 0.00998954 | 0.10894399 | 0.09169431 | 0.92694091 | -0.2035368 | 0.22351584 | PYBI022 | model5 | 1556.78833 |
| dist_aq.ag | 2.55767237 | 40.5899632 | 0.06301243 | 0.9497566 | -76.997194 | 82.1125383 | PYBI022 | model6 | 1566.61651 |
| log_sl | -53.143397 | 67.1060658 | -0.7919313 | 0.42840071 | -184.66887 | 78.3820749 | PYBI022 | model6 | 1566.61651 |
| cos_ta | -180.7729 | 116.350844 | -1.5536879 | 0.12025889 | -408.81636 | 47.2705682 | PYBI022 | model6 | 1566.61651 |
| dist_aq.ag:log_sl | 5.55439231 | 7.01260369 | 0.79205849 | 0.42832657 | -8.1900583 | 19.298843 | PYBI022 | model6 | 1566.61651 |
| dist_aq.ag:cos_ta | 18.886416 | 12.1597147 | 1.55319565 | 0.1203764 | -4.9461869 | 42.7190189 | PYBI022 | model6 | 1566.61651 |
| log_sl:cos_ta | -0.0085173 | 0.10669737 | -0.0798265 | 0.93637522 | -0.2176403 | 0.20060572 | PYBI022 | model6 | 1566.61651 |
| dist_terr.ag | 15.9823863 | 12.193737 | 1.31070453 | 0.18995761 | -7.916899 | 39.8816716 | PYBI022 | model7 | 1565.75723 |
| log_sl | 15.3763624 | 18.5726607 | 0.82790305 | 0.40772541 | -21.025384 | 51.7781084 | PYBI022 | model7 | 1565.75723 |
| cos_ta | 55.267858 | 30.7274383 | 1.79864841 | 0.07207431 | -4.9568144 | 115.49253 | PYBI022 | model7 | 1565.75723 |

| | | | | | | | | | |
|---|---|---|---|---|---|---|---|---|---|
| dist_terr.ag:log_sl | -1.869343 | 2.2589953 | -0.8275108 | 0.40794761 | -6.2968924 | 2.55820644 | PYBI022 | model7 | 1565.75723 |
| dist_terr.ag:cos_ta | -6.7239728 | 3.73382823 | -1.8008254 | 0.0717304 | -14.042142 | 0.5941961 | PYBI022 | model7 | 1565.75723 |
| log_sl:cos_ta | -0.0123328 | 0.10726163 | -0.1149788 | 0.90846189 | -0.2225617 | 0.19789611 | PYBI022 | model7 | 1565.75723 |
| dist_road | 6.97147169 | 4.16174698 | 1.67513107 | 0.09390852 | -1.1854025 | 15.1283459 | PYBI022 | model8 | 1567.82586 |
| dist_forest | -2.1567274 | 2.45558063 | -0.8782963 | 0.37978294 | -6.969577 | 2.6561222 | PYBI022 | model8 | 1567.82586 |
| dist_settle | 5.08816983 | 11.9496298 | 0.42580146 | 0.67025252 | -18.332674 | 28.5090138 | PYBI022 | model8 | 1567.82586 |
| log_sl | 0.01775669 | 0.07569174 | 0.23459218 | 0.81452528 | -0.1305964 | 0.16610977 | PYBI022 | model8 | 1567.82586 |
| cos_ta | -0.0568138 | 0.4397881 | -0.1291844 | 0.89721174 | -0.9187826 | 0.80515508 | PYBI022 | model8 | 1567.82586 |
| log_sl:cos_ta | -0.001069 | 0.10650322 | -0.0100372 | 0.99199158 | -0.2098115 | 0.20767349 | PYBI022 | model8 | 1567.82586 |
| dist_road | 2.81290706 | 4.2663879 | 0.65931817 | 0.50969148 | -5.5490596 | 11.1748737 | PYBI022 | model9 | 1563.61566 |
| dist_terr.ag | -1.4331337 | 6.04028788 | -0.2372625 | 0.81245316 | -13.27188 | 10.405613 | PYBI022 | model9 | 1563.61566 |
| dist_water | 32.4909717 | 15.2093179 | 2.13625436 | 0.03265869 | 2.68125639 | 62.300687 | PYBI022 | model9 | 1563.61566 |
| log_sl | 0.02419953 | 0.07581295 | 0.31920052 | 0.74957446 | -0.1243911 | 0.17279018 | PYBI022 | model9 | 1563.61566 |
| cos_ta | -0.0711871 | 0.44015493 | -0.1617319 | 0.87151701 | -0.9338749 | 0.79150073 | PYBI022 | model9 | 1563.61566 |
| log_sl:cos_ta | 0.00619522 | 0.10670131 | 0.05806134 | 0.95369977 | -0.2029355 | 0.21532594 | PYBI022 | model9 | 1563.61566 |
| dist_water | 34.7580359 | 13.1328867 | 2.64664096 | 0.00812956 | 9.01805092 | 60.4980209 | PYBI022 | model10 | 1562.47217 |
| dist_settle | 3.30998176 | 9.18152602 | 0.36050453 | 0.71846987 | -14.685479 | 21.3054421 | PYBI022 | model10 | 1562.47217 |
| dist_aq.ag | 32.513844 | 25.8226387 | 1.25912167 | 0.20798639 | -18.097598 | 83.1252859 | PYBI022 | model10 | 1562.47217 |
| log_sl | 0.02792761 | 0.07601582 | 0.3673921 | 0.71332656 | -0.1210607 | 0.17691587 | PYBI022 | model10 | 1562.47217 |
| cos_ta | -0.0752928 | 0.44154144 | -0.1705225 | 0.86459921 | -0.9406981 | 0.79011256 | PYBI022 | model10 | 1562.47217 |
| log_sl:cos_ta | 0.00757865 | 0.10707138 | 0.07078129 | 0.94357182 | -0.2022774 | 0.21743469 | PYBI022 | model10 | 1562.47217 |
| log_sl | -0.0209236 | 0.08548992 | -0.2447492 | 0.80665062 | -0.1884808 | 0.14663357 | PYBI028 | model1 | 725.440246 |
| cos_ta | -0.3120185 | 0.56937136 | -0.5480052 | 0.58368832 | -1.4279658 | 0.80392888 | PYBI028 | model1 | 725.440246 |
| log_sl:cos_ta | 0.01718151 | 0.12126296 | 0.14168802 | 0.88732644 | -0.2204895 | 0.25485254 | PYBI028 | model1 | 725.440246 |
| dist_forest | -13.14144 | 6.95386619 | -1.8898034 | 0.05878425 | -26.770767 | 0.48788711 | PYBI028 | model2 | 721.248766 |
| log_sl | -23.745182 | 8.91870276 | -2.6624031 | 0.00775849 | -41.225518 | -6.2648454 | PYBI028 | model2 | 721.248766 |
| cos_ta | 18.5859523 | 18.4651665 | 1.00654128 | 0.31415527 | -17.605109 | 54.7770136 | PYBI028 | model2 | 721.248766 |
| dist_forest:log_sl | 3.05018882 | 1.14831053 | 2.65624041 | 0.00790173 | 0.79954154 | 5.3008361 | PYBI028 | model2 | 721.248766 |
| dist_forest:cos_ta | -2.4412991 | 2.38751364 | -1.0225278 | 0.30653116 | -7.1207398 | 2.23814166 | PYBI028 | model2 | 721.248766 |
| log_sl:cos_ta | 0.04930942 | 0.11976433 | 0.41172041 | 0.68054436 | -0.1854243 | 0.28404319 | PYBI028 | model2 | 721.248766 |
| dist_settle | 99.9382192 | 36.1314493 | 2.76596209 | 0.00567551 | 29.1218798 | 170.754559 | PYBI028 | model3 | 720.70499 |
| log_sl | 142.534304 | 50.5672185 | 2.81870959 | 0.00482171 | 43.4243766 | 241.644231 | PYBI028 | model3 | 720.70499 |
| cos_ta | 58.5964644 | 76.5874052 | 0.7650927 | 0.44421635 | -91.512092 | 208.70502 | PYBI028 | model3 | 720.70499 |
| dist_settle:log_sl | -16.057711 | 5.69482449 | -2.8197025 | 0.00480682 | -27.219362 | -4.8960597 | PYBI028 | model3 | 720.70499 |
| dist_settle:cos_ta | -6.6230256 | 8.61103179 | -0.7691326 | 0.44181458 | -23.500338 | 10.2542866 | PYBI028 | model3 | 720.70499 |
| log_sl:cos_ta | -0.0113272 | 0.11981038 | -0.0945425 | 0.92467826 | -0.2461512 | 0.22349687 | PYBI028 | model3 | 720.70499 |
| dist_road | 34.3933271 | 17.1614759 | 2.00410077 | 0.04505927 | 0.75745237 | 68.0292018 | PYBI028 | model4 | 726.273322 |
| log_sl | 44.0039309 | 21.2785995 | 2.06798999 | 0.03864096 | 2.29864221 | 85.7092196 | PYBI028 | model4 | 726.273322 |
| cos_ta | -28.002567 | 40.2754152 | -0.6952769 | 0.48688177 | -106.94093 | 50.9357962 | PYBI028 | model4 | 726.273322 |
| dist_road:log_sl | -5.6118473 | 2.71134717 | -2.0697635 | 0.0384745 | -10.92599 | -0.2977045 | PYBI028 | model4 | 726.273322 |

| | | | | | | | | | |
|---|---|---|---|---|---|---|---|---|---|
| dist_road:cos_ta | 3.51614179 | 5.11885381 | 0.68690022 | 0.49214561 | -6.5166273 | 13.5489109 | PYBI028 | model4 | 726.273322 |
| log_sl:cos_ta | 0.04174649 | 0.12133513 | 0.34405933 | 0.73080167 | -0.196066 | 0.27955898 | PYBI028 | model4 | 726.273322 |
| dist_water | 19.6667045 | 16.4826121 | 1.19317887 | 0.23279933 | -12.638622 | 51.9720306 | PYBI028 | model5 | 718.981581 |
| log_sl | 48.4360855 | 20.93623 | 2.3135056 | 0.02069486 | 7.40182865 | 89.4703423 | PYBI028 | model5 | 718.981581 |
| cos_ta | -27.860955 | 32.3607593 | -0.8609487 | 0.38926628 | -91.286877 | 35.5649681 | PYBI028 | model5 | 718.981581 |
| dist_water:log_sl | -5.5240806 | 2.38664967 | -2.3145754 | 0.02063618 | -10.201828 | -0.8463332 | PYBI028 | model5 | 718.981581 |
| dist_water:cos_ta | 3.14025629 | 3.67697543 | 0.85403244 | 0.39308702 | -4.0664831 | 10.3469957 | PYBI028 | model5 | 718.981581 |
| log_sl:cos_ta | 0.02151365 | 0.12033992 | 0.17877398 | 0.85811518 | -0.2143483 | 0.25737555 | PYBI028 | model5 | 718.981581 |
| dist_aq.ag | 82.2290043 | 26.900191 | 3.0568186 | 0.002237 | 29.5055987 | 134.95241 | PYBI028 | model6 | 713.657447 |
| log_sl | 133.237688 | 35.6278308 | 3.73970811 | 1.84E-04 | 63.4084227 | 203.066953 | PYBI028 | model6 | 713.657447 |
| cos_ta | -29.990319 | 52.9679952 | -0.566197 | 0.57125988 | -133.80568 | 73.8250438 | PYBI028 | model6 | 713.657447 |
| dist_aq.ag:log_sl | -13.966832 | 3.73270131 | -3.7417491 | 1.83E-04 | -21.282792 | -6.6508716 | PYBI028 | model6 | 713.657447 |
| dist_aq.ag:cos_ta | 3.11036363 | 5.53606838 | 0.5618362 | 0.57422762 | -7.740131 | 13.9608583 | PYBI028 | model6 | 713.657447 |
| log_sl:cos_ta | 0.01844643 | 0.11464156 | 0.1609053 | 0.87216799 | -0.2062469 | 0.24313976 | PYBI028 | model6 | 713.657447 |
| dist_terr.ag | 53.1803039 | 23.7385976 | 2.24024624 | 0.02507494 | 6.65350762 | 99.7071001 | PYBI028 | model7 | 724.979727 |
| log_sl | 66.3148964 | 29.8748174 | 2.21975905 | 0.02643513 | 7.76133014 | 124.868463 | PYBI028 | model7 | 724.979727 |
| cos_ta | -39.483249 | 49.2259405 | -0.8020822 | 0.42250543 | -135.96432 | 56.9978219 | PYBI028 | model7 | 724.979727 |
| dist_terr.ag:log_sl | -8.0481852 | 3.62363328 | -2.2210264 | 0.02634917 | -15.150376 | -0.9459944 | PYBI028 | model7 | 724.979727 |
| dist_terr.ag:cos_ta | 4.74731675 | 5.96135627 | 0.79634844 | 0.42582954 | -6.9367268 | 16.4313603 | PYBI028 | model7 | 724.979727 |
| log_sl:cos_ta | 0.0374485 | 0.11928278 | 0.31394724 | 0.75356111 | -0.1963415 | 0.27123846 | PYBI028 | model7 | 724.979727 |
| dist_road | 1.23337189 | 4.74441909 | 0.25996268 | 0.79489256 | -8.0655187 | 10.5322624 | PYBI028 | model8 | 728.493989 |
| dist_forest | 3.54031349 | 2.36474395 | 1.49712339 | 0.13436115 | -1.0944995 | 8.17512646 | PYBI028 | model8 | 728.493989 |
| dist_settle | 7.47800017 | 12.0447532 | 0.62085126 | 0.53469749 | -16.129282 | 31.0852826 | PYBI028 | model8 | 728.493989 |
| log_sl | -0.0042195 | 0.08768162 | -0.0481227 | 0.96161845 | -0.1760723 | 0.16763334 | PYBI028 | model8 | 728.493989 |
| cos_ta | -0.333216 | 0.58029812 | -0.5742152 | 0.56582219 | -1.4705794 | 0.80414742 | PYBI028 | model8 | 728.493989 |
| log_sl:cos_ta | 0.02525474 | 0.12407921 | 0.20353725 | 0.83871514 | -0.217936 | 0.26844553 | PYBI028 | model8 | 728.493989 |
| dist_road | 2.57895032 | 5.15864555 | 0.4999278 | 0.61712592 | -7.5318092 | 12.6897098 | PYBI028 | model9 | 717.278333 |
| dist_terr.ag | 26.1027664 | 9.96278326 | 2.62002753 | 0.00879227 | 6.57607001 | 45.6294628 | PYBI028 | model9 | 717.278333 |
| dist_water | -28.697655 | 7.94456832 | -3.6122359 | 3.04E-04 | -44.268723 | -13.126587 | PYBI028 | model9 | 717.278333 |
| log_sl | 0.04935492 | 0.09102987 | 0.54218383 | 0.58769187 | -0.1290603 | 0.22777019 | PYBI028 | model9 | 717.278333 |
| cos_ta | -0.3536249 | 0.59241529 | -0.5969206 | 0.55056037 | -1.5147375 | 0.80748772 | PYBI028 | model9 | 717.278333 |
| log_sl:cos_ta | 0.04357963 | 0.12726057 | 0.3424441 | 0.7320167 | -0.2058465 | 0.29300578 | PYBI028 | model9 | 717.278333 |
| dist_water | -18.85846 | 7.62817639 | -2.4722108 | 0.01342803 | -33.809411 | -3.9075092 | PYBI028 | model10 | 723.75729 |
| dist_settle | 13.0586554 | 11.528857 | 1.13269298 | 0.25734321 | -9.5374891 | 35.6547999 | PYBI028 | model10 | 723.75729 |
| dist_aq.ag | 5.70296338 | 13.6806578 | 0.41686324 | 0.67677844 | -21.110633 | 32.51656 | PYBI028 | model10 | 723.75729 |
| log_sl | 0.01945757 | 0.08931862 | 0.21784451 | 0.82755026 | -0.1556037 | 0.19451885 | PYBI028 | model10 | 723.75729 |
| cos_ta | -0.2812536 | 0.58492846 | -0.4808342 | 0.63063436 | -1.4276923 | 0.86518512 | PYBI028 | model10 | 723.75729 |
| log_sl:cos_ta | 0.0119453 | 0.12496135 | 0.095592 | 0.92384462 | -0.2329744 | 0.25686504 | PYBI028 | model10 | 723.75729 |
| log_sl | -0.0087511 | 0.05400022 | -0.1620575 | 0.87126058 | -0.1145896 | 0.09708734 | PYBI029 | model1 | 2456.00099 |
| cos_ta | -0.1146397 | 0.33001649 | -0.3473756 | 0.72830919 | -0.7614601 | 0.53218078 | PYBI029 | model1 | 2456.00099 |

| | | | | | | | | | |
|---|---|---|---|---|---|---|---|---|---|
| log_sl:cos_ta | 0.02653081 | 0.07566822 | 0.35062025 | 0.72587326 | -0.1217762 | 0.1748378 | PYBI029 | model1 | 2456.00099 |
| dist_forest | 5.20554268 | 2.32399778 | 2.23990863 | 0.02509686 | 0.65059073 | 9.76049463 | PYBI029 | model2 | 2452.2252 |
| log_sl | 2.31726293 | 2.82021334 | 0.82166228 | 0.41126913 | -3.2102536 | 7.8447795 | PYBI029 | model2 | 2452.2252 |
| cos_ta | -3.7026587 | 5.21758597 | -0.7096498 | 0.47792134 | -13.928939 | 6.52362188 | PYBI029 | model2 | 2452.2252 |
| dist_forest:log_sl | -0.3014218 | 0.36691244 | -0.8215088 | 0.41135651 | -1.020557 | 0.41771336 | PYBI029 | model2 | 2452.2252 |
| dist_forest:cos_ta | 0.46064961 | 0.67587108 | 0.68156432 | 0.49551448 | -0.8640334 | 1.78533259 | PYBI029 | model2 | 2452.2252 |
| log_sl:cos_ta | 0.04526736 | 0.07651287 | 0.59163066 | 0.55409794 | -0.1046951 | 0.19522983 | PYBI029 | model2 | 2452.2252 |
| dist_settle | -10.311896 | 9.67963631 | -1.0653185 | 0.28673183 | -29.283634 | 8.65984296 | PYBI029 | model3 | 2458.8896 |
| log_sl | -7.2078649 | 14.5255025 | -0.4962214 | 0.61973822 | -35.677327 | 21.2615967 | PYBI029 | model3 | 2458.8896 |
| cos_ta | -35.158279 | 27.7807403 | -1.2655631 | 0.20566955 | -89.607529 | 19.2909712 | PYBI029 | model3 | 2458.8896 |
| dist_settle:log_sl | 0.82168935 | 1.65720339 | 0.49582891 | 0.62001513 | -2.4263696 | 4.06974831 | PYBI029 | model3 | 2458.8896 |
| dist_settle:cos_ta | 3.99905445 | 3.17065275 | 1.26127166 | 0.20721099 | -2.2153108 | 10.2134196 | PYBI029 | model3 | 2458.8896 |
| log_sl:cos_ta | 0.02322476 | 0.076174 | 0.30489088 | 0.76044926 | -0.1260735 | 0.17252305 | PYBI029 | model3 | 2458.8896 |
| dist_road | -6.2609726 | 6.79360583 | -0.9215979 | 0.35673838 | -19.576195 | 7.05425019 | PYBI029 | model4 | 2456.14173 |
| log_sl | -17.176373 | 9.83303622 | -1.7468026 | 0.08067159 | -36.44877 | 2.09602388 | PYBI029 | model4 | 2456.14173 |
| cos_ta | 7.58900599 | 19.9394448 | 0.38060267 | 0.7034981 | -31.491588 | 46.6695996 | PYBI029 | model4 | 2456.14173 |
| dist_road:log_sl | 2.19260777 | 1.25556411 | 1.74631288 | 0.0807566 | -0.2682527 | 4.6534682 | PYBI029 | model4 | 2456.14173 |
| dist_road:cos_ta | -0.9837735 | 2.54750561 | -0.3861713 | 0.69936982 | -5.9767927 | 4.00924575 | PYBI029 | model4 | 2456.14173 |
| log_sl:cos_ta | 0.0279419 | 0.07618906 | 0.36674424 | 0.71380981 | -0.1213859 | 0.1772697 | PYBI029 | model4 | 2456.14173 |
| dist_water | 7.57884664 | 23.6030915 | 0.32109551 | 0.74813801 | -38.682363 | 53.8400559 | PYBI029 | model5 | 2438.10153 |
| log_sl | -62.474843 | 43.5024033 | -1.436124 | 0.15096707 | -147.73799 | 22.7883004 | PYBI029 | model5 | 2438.10153 |
| cos_ta | 77.226602 | 87.6561489 | 0.88101751 | 0.37830834 | -94.576293 | 249.029497 | PYBI029 | model5 | 2438.10153 |
| dist_water:log_sl | 7.08919725 | 4.93474983 | 1.43658696 | 0.15083539 | -2.5827347 | 16.7611292 | PYBI029 | model5 | 2438.10153 |
| dist_water:cos_ta | -8.7740604 | 9.94147124 | -0.8825716 | 0.37746776 | -28.258986 | 10.7108651 | PYBI029 | model5 | 2438.10153 |
| log_sl:cos_ta | 0.0344535 | 0.0773436 | 0.44546021 | 0.65598721 | -0.1171372 | 0.18604416 | PYBI029 | model5 | 2438.10153 |
| dist_aq.ag | 73.9615925 | 127.398309 | 0.58055396 | 0.56154111 | -175.73451 | 323.65769 | PYBI029 | model6 | 2460.39387 |
| log_sl | 70.9554158 | 240.851425 | 0.29460243 | 0.7682976 | -401.1047 | 543.015535 | PYBI029 | model6 | 2460.39387 |
| cos_ta | -149.09173 | 441.485897 | -0.3377044 | 0.73558596 | #NAME? | Inf | PYBI029 | model6 | 2460.39387 |
| dist_aq.ag:log_sl | -7.411162 | 25.1541833 | -0.2946294 | 0.768277 | -56.712455 | 41.8901314 | PYBI029 | model6 | 2460.39387 |
| dist_aq.ag:cos_ta | 15.5585616 | 46.1069415 | 0.3374451 | 0.73578139 | -74.809383 | 105.926506 | PYBI029 | model6 | 2460.39387 |
| log_sl:cos_ta | 0.026992 | 0.07584855 | 0.35586706 | 0.72194012 | -0.1216684 | 0.17565242 | PYBI029 | model6 | 2460.39387 |
| dist_terr.ag | -12.937223 | 10.0539277 | -1.286783 | 0.19816993 | -32.64256 | 6.76811285 | PYBI029 | model7 | 2453.68405 |
| log_sl | -33.903992 | 15.4207964 | -2.1985889 | 0.02790717 | -64.128197 | -3.6797861 | PYBI029 | model7 | 2453.68405 |
| cos_ta | -0.9600174 | 30.7558044 | -0.0312142 | 0.97509873 | -61.240286 | 59.3202516 | PYBI029 | model7 | 2453.68405 |
| dist_terr.ag:log_sl | 4.12101839 | 1.87491621 | 2.19797469 | 0.02795091 | 0.44625013 | 7.79578664 | PYBI029 | model7 | 2453.68405 |
| dist_terr.ag:cos_ta | 0.10240193 | 3.74144751 | 0.0273696 | 0.97816494 | -7.2307004 | 7.4355043 | PYBI029 | model7 | 2453.68405 |
| log_sl:cos_ta | 0.02843651 | 0.07614626 | 0.37344588 | 0.70881662 | -0.1208074 | 0.17768043 | PYBI029 | model7 | 2453.68405 |
| dist_road | 3.0934998 | 2.79986791 | 1.10487348 | 0.26921442 | -2.3941405 | 8.58114007 | PYBI029 | model8 | 2444.86425 |
| dist_forest | 5.60014727 | 1.59959253 | 3.50098363 | 4.64E-04 | 2.46500352 | 8.73529102 | PYBI029 | model8 | 2444.86425 |
| dist_settle | -13.620161 | 5.36282648 | -2.5397355 | 0.01109363 | -24.131108 | -3.1092142 | PYBI029 | model8 | 2444.86425 |

| | | | | | | | | | |
|---|---|---|---|---|---|---|---|---|---|
| log_sl | 0.02206751 | 0.05566456 | 0.39643734 | 0.69178243 | -0.087033 | 0.13116803 | PYBI029 | model8 | 2444.86425 |
| cos_ta | -0.1870379 | 0.33929673 | -0.5512518 | 0.58146108 | -0.8520473 | 0.47797144 | PYBI029 | model8 | 2444.86425 |
| log_sl:cos_ta | 0.05185503 | 0.07831183 | 0.66216091 | 0.5078681 | -0.1016333 | 0.2053434 | PYBI029 | model8 | 2444.86425 |
| dist_road | 2.88400082 | 2.98867699 | 0.96497575 | 0.33455696 | -2.9736984 | 8.74170008 | PYBI029 | model9 | 2439.77213 |
| dist_terr.ag | -1.8116594 | 4.96323709 | -0.3650157 | 0.7150997 | -11.539425 | 7.91610658 | PYBI029 | model9 | 2439.77213 |
| dist_water | 36.9302665 | 8.8026761 | 4.19534538 | 2.72E-05 | 19.6773384 | 54.1831946 | PYBI029 | model9 | 2439.77213 |
| log_sl | 0.02671521 | 0.0557138 | 0.47950798 | 0.63157729 | -0.0824818 | 0.13591226 | PYBI029 | model9 | 2439.77213 |
| cos_ta | -0.1387167 | 0.33670905 | -0.4119779 | 0.68035561 | -0.7986543 | 0.52122092 | PYBI029 | model9 | 2439.77213 |
| log_sl:cos_ta | 0.03650899 | 0.07737083 | 0.47187017 | 0.63701946 | -0.1151351 | 0.18815303 | PYBI029 | model9 | 2439.77213 |
| dist_water | 37.3677028 | 8.26678011 | 4.5202246 | 6.18E-06 | 21.1651115 | 53.5702941 | PYBI029 | model10 | 2437.96269 |
| dist_settle | -6.6292859 | 4.5756286 | -1.4488252 | 0.14738641 | -15.597353 | 2.33878135 | PYBI029 | model10 | 2437.96269 |
| dist_aq.ag | 30.7522055 | 37.8806893 | 0.81181747 | 0.41689637 | -43.492581 | 104.996992 | PYBI029 | model10 | 2437.96269 |
| log_sl | 0.02993998 | 0.05584071 | 0.53616753 | 0.59184277 | -0.0795058 | 0.13938576 | PYBI029 | model10 | 2437.96269 |
| cos_ta | -0.1341825 | 0.33717225 | -0.3979644 | 0.69065644 | -0.795028 | 0.52666292 | PYBI029 | model10 | 2437.96269 |
| log_sl:cos_ta | 0.0345758 | 0.07744502 | 0.44645606 | 0.65526785 | -0.1172136 | 0.18636524 | PYBI029 | model10 | 2437.96269 |
| log_sl | -0.0229963 | 0.21801209 | -0.1054816 | 0.91599365 | -0.4502921 | 0.40429958 | PYBI033 | model1 | 228.63292 |
| cos_ta | -0.3388679 | 1.21385158 | -0.2791675 | 0.78011629 | -2.7179733 | 2.04023748 | PYBI033 | model1 | 228.63292 |
| log_sl:cos_ta | 0.09685293 | 0.30980569 | 0.31262475 | 0.75456577 | -0.5103551 | 0.70406093 | PYBI033 | model1 | 228.63292 |
| dist_forest | -4.2820497 | 16.3488177 | -0.261918 | 0.79338466 | -36.325144 | 27.7610442 | PYBI033 | model2 | 232.58281 |
| log_sl | -12.016826 | 21.8179407 | -0.5507773 | 0.58178636 | -54.779204 | 30.7455521 | PYBI033 | model2 | 232.58281 |
| cos_ta | 44.3215748 | 33.3249605 | 1.32998132 | 0.18352443 | -20.994148 | 109.637297 | PYBI033 | model2 | 232.58281 |
| dist_forest:log_sl | 1.55339527 | 2.82869664 | 0.54915584 | 0.58289851 | -3.9907483 | 7.09753881 | PYBI033 | model2 | 232.58281 |
| dist_forest:cos_ta | -5.8068517 | 4.32791818 | -1.3417194 | 0.179687 | -14.289415 | 2.67571202 | PYBI033 | model2 | 232.58281 |
| log_sl:cos_ta | 0.14408659 | 0.3218666 | 0.44765933 | 0.65439908 | -0.4867604 | 0.77493353 | PYBI033 | model2 | 232.58281 |
| dist_settle | -117.26178 | 103.157716 | -1.1367233 | 0.25565399 | -319.44718 | 84.9236334 | PYBI033 | model3 | 228.086243 |
| log_sl | -357.97581 | 204.568062 | -1.7499106 | 0.08013375 | #NAME? | 42.9702216 | PYBI033 | model3 | 228.086243 |
| cos_ta | -555.76324 | 380.028932 | -1.4624235 | 0.14362519 | #NAME? | 189.079778 | PYBI033 | model3 | 228.086243 |
| dist_settle:log_sl | 40.3236372 | 23.0481256 | 1.74954084 | 0.08019758 | -4.849859 | 85.4971334 | PYBI033 | model3 | 228.086243 |
| dist_settle:cos_ta | 62.6198308 | 42.8531023 | 1.46126715 | 0.14394214 | -21.370706 | 146.610368 | PYBI033 | model3 | 228.086243 |
| log_sl:cos_ta | -0.0522855 | 0.32354021 | -0.1616043 | 0.87161744 | -0.6864127 | 0.58184166 | PYBI033 | model3 | 228.086243 |
| dist_road | -20.472192 | 36.3084358 | -0.5638412 | 0.57286222 | -91.635418 | 50.6910349 | PYBI033 | model4 | 227.849348 |
| log_sl | -74.731384 | 66.7851553 | -1.118982 | 0.26314783 | -205.62788 | 56.165115 | PYBI033 | model4 | 227.849348 |
| cos_ta | -194.64148 | 105.339135 | -1.8477603 | 0.06463702 | -401.10239 | 11.8194349 | PYBI033 | model4 | 227.849348 |
| dist_road:log_sl | 9.52675324 | 8.51808475 | 1.11841494 | 0.26338982 | -7.1683861 | 26.2218926 | PYBI033 | model4 | 227.849348 |
| dist_road:cos_ta | 24.8168885 | 13.4555193 | 1.84436498 | 0.06512997 | -1.5554446 | 51.1892217 | PYBI033 | model4 | 227.849348 |
| log_sl:cos_ta | -0.0149157 | 0.33428322 | -0.0446198 | 0.96441034 | -0.6700987 | 0.64026741 | PYBI033 | model4 | 227.849348 |
| dist_water | 31.8768386 | 171.78583 | 0.18556151 | 0.85278864 | -304.8172 | 368.570879 | PYBI033 | model5 | 228.754028 |
| log_sl | -22.566726 | 340.175508 | -0.0663385 | 0.94710835 | -689.29847 | 644.165019 | PYBI033 | model5 | 228.754028 |
| cos_ta | -976.07864 | 480.035967 | -2.0333448 | 0.0420177 | #NAME? | -35.225428 | PYBI033 | model5 | 228.754028 |
| dist_water:log_sl | 2.56055821 | 38.562426 | 0.06640034 | 0.9470591 | -73.020408 | 78.1415243 | PYBI033 | model5 | 228.754028 |

| | | | | | | | | | |
|---|---|---|---|---|---|---|---|---|---|
| dist_water:cos_ta | 110.586939 | 54.4085405 | 2.03252905 | 0.04210013 | 3.94815948 | 217.225719 | PYBI033 | model5 | 228.754028 |
| log_sl:cos_ta | 0.16228182 | 0.33641208 | 0.48238998 | 0.62952894 | -0.4970737 | 0.82163737 | PYBI033 | model5 | 228.754028 |
| dist_aq.ag | 10.3301693 | 123.044056 | 0.08395504 | 0.93309217 | -230.83175 | 251.492088 | PYBI033 | model6 | 227.177794 |
| log_sl | 334.281679 | 219.151886 | 1.52534246 | 0.1271737 | -95.248126 | Inf | PYBI033 | model6 | 227.177794 |
| cos_ta | -547.55062 | 373.966334 | -1.4641709 | 0.14314726 | #NAME? | 185.409925 | PYBI033 | model6 | 227.177794 |
| dist_aq.ag:log_sl | -35.056521 | 22.9786377 | -1.5256135 | 0.12710614 | -80.093823 | 9.98078141 | PYBI033 | model6 | 227.177794 |
| dist_aq.ag:cos_ta | 57.311756 | 39.1919598 | 1.46233453 | 0.14364956 | -19.503074 | 134.126586 | PYBI033 | model6 | 227.177794 |
| log_sl:cos_ta | 0.36848326 | 0.32227066 | 1.14339685 | 0.25287386 | -0.2631556 | 1.00012215 | PYBI033 | model6 | 227.177794 |
| dist_terr.ag | -70.26502 | 188.214191 | -0.3733248 | 0.70890674 | -439.15806 | 298.628016 | PYBI033 | model7 | 230.81822 |
| log_sl | -321.60062 | 366.018224 | -0.8786465 | 0.379593 | #NAME? | 395.781919 | PYBI033 | model7 | 230.81822 |
| cos_ta | 768.731982 | 611.790979 | 1.25652716 | 0.2089249 | -430.3563 | Inf | PYBI033 | model7 | 230.81822 |
| dist_terr.ag:log_sl | 38.9821701 | 44.3726063 | 0.87851883 | 0.37966222 | -47.98654 | 125.95088 | PYBI033 | model7 | 230.81822 |
| dist_terr.ag:cos_ta | -93.261975 | 74.1880809 | -1.2571019 | 0.20871675 | -238.66794 | 52.1439913 | PYBI033 | model7 | 230.81822 |
| log_sl:cos_ta | 0.18729939 | 0.31518815 | 0.5942463 | 0.55234739 | -0.430458 | 0.80505681 | PYBI033 | model7 | 230.81822 |
| dist_road | 15.629678 | 19.7877856 | 0.78986494 | 0.42960665 | -23.153669 | 54.4130252 | PYBI033 | model8 | 232.650963 |
| dist_forest | 3.33018331 | 9.06599582 | 0.36732681 | 0.71337526 | -14.438842 | 21.0992086 | PYBI033 | model8 | 232.650963 |
| dist_settle | 15.095733 | 61.5880184 | 0.24510828 | 0.80637259 | -105.61456 | 135.806031 | PYBI033 | model8 | 232.650963 |
| log_sl | -0.0235624 | 0.21965933 | -0.1072679 | 0.91457644 | -0.4540868 | 0.40696198 | PYBI033 | model8 | 232.650963 |
| cos_ta | -0.3816629 | 1.20931255 | -0.3156032 | 0.75230371 | -2.7518719 | 1.98854614 | PYBI033 | model8 | 232.650963 |
| log_sl:cos_ta | 0.11668068 | 0.31132203 | 0.37479095 | 0.70781595 | -0.4934993 | 0.72686064 | PYBI033 | model8 | 232.650963 |
| dist_road | 27.8685301 | 14.2284833 | 1.95864377 | 0.05015452 | -0.0187847 | 55.7558449 | PYBI033 | model9 | 227.458143 |
| dist_terr.ag | 101.67941 | 56.24233 | 1.80788047 | 0.07062511 | -8.553531 | 211.912351 | PYBI033 | model9 | 227.458143 |
| dist_water | 61.2903038 | 46.326995 | 1.3229933 | 0.18583761 | -29.508938 | 152.089545 | PYBI033 | model9 | 227.458143 |
| log_sl | 0.02960912 | 0.22714847 | 0.13035138 | 0.89628843 | -0.4155937 | 0.47481194 | PYBI033 | model9 | 227.458143 |
| cos_ta | -0.2463053 | 1.23181263 | -0.1999536 | 0.8415169 | -2.6606137 | 2.16800307 | PYBI033 | model9 | 227.458143 |
| log_sl:cos_ta | 0.11040699 | 0.31564546 | 0.34978165 | 0.72650257 | -0.5082467 | 0.72906072 | PYBI033 | model9 | 227.458143 |
| dist_water | 105.485013 | 54.7558557 | 1.92646086 | 0.05404686 | -1.8344925 | 212.804518 | PYBI033 | model10 | 226.236962 |
| dist_settle | 48.9340782 | 46.3008987 | 1.05687102 | 0.29057046 | -41.814016 | 139.682172 | PYBI033 | model10 | 226.236962 |
| dist_aq.ag | -155.44382 | 73.9984563 | -2.100636 | 0.03567293 | -300.47813 | -10.40951 | PYBI033 | model10 | 226.236962 |
| log_sl | 0.06900916 | 0.23030808 | 0.29963845 | 0.76445295 | -0.3823864 | 0.5204047 | PYBI033 | model10 | 226.236962 |
| cos_ta | -0.8481072 | 1.26030058 | -0.6729404 | 0.50098522 | -3.3182509 | 1.62203658 | PYBI033 | model10 | 226.236962 |
| log_sl:cos_ta | 0.32672205 | 0.33414175 | 0.97779475 | 0.32817585 | -0.3281837 | 0.98162784 | PYBI033 | model10 | 226.236962 |
| log_sl | -0.0407311 | 0.12555807 | -0.3244005 | 0.74563487 | -0.2868204 | 0.2053582 | PYBI055 | model1 | 269.387671 |
| cos_ta | -0.9878936 | 0.74426569 | -1.32734 | 0.18439624 | -2.4466276 | 0.47084034 | PYBI055 | model1 | 269.387671 |
| log_sl:cos_ta | 0.22433114 | 0.17344246 | 1.29340378 | 0.19587144 | -0.1156098 | 0.56427212 | PYBI055 | model1 | 269.387671 |
| dist_forest | 11.4855746 | 12.4871034 | 0.91979495 | 0.35767992 | -12.988698 | 35.9598474 | PYBI055 | model2 | 273.901152 |
| log_sl | 9.40481603 | 17.7227827 | 0.53066249 | 0.59565268 | -25.3312 | 44.1408319 | PYBI055 | model2 | 273.901152 |
| cos_ta | 11.6693923 | 36.596361 | 0.31886756 | 0.74982695 | -60.058157 | 83.3969419 | PYBI055 | model2 | 273.901152 |
| dist_forest:log_sl | -1.2101226 | 2.26909132 | -0.5333071 | 0.59382099 | -5.6574599 | 3.23721468 | PYBI055 | model2 | 273.901152 |
| dist_forest:cos_ta | -1.6163304 | 4.68075409 | -0.3453141 | 0.72985824 | -10.79044 | 7.55777902 | PYBI055 | model2 | 273.901152 |

| | | | | | | | | | |
|---|---|---|---|---|---|---|---|---|---|
| log_sl:cos_ta | 0.21609188 | 0.173191 | 1.24770853 | 0.21213781 | -0.1233562 | 0.55554 | PYBI055 | model2 | 273.901152 |
| dist_settle | 68.8951997 | 51.3658166 | 1.34126554 | 0.17983425 | -31.779951 | 169.57035 | PYBI055 | model3 | 269.769705 |
| log_sl | 73.7023551 | 83.1878529 | 0.88597497 | 0.37563102 | -89.342841 | 236.747551 | PYBI055 | model3 | 269.769705 |
| cos_ta | 334.341424 | 191.659763 | 1.74445287 | 0.08108015 | -41.30481 | Inf | PYBI055 | model3 | 269.769705 |
| dist_settle:log_sl | -8.3079573 | 9.37237905 | -0.8864299 | 0.3753859 | -26.677483 | 10.0615681 | PYBI055 | model3 | 269.769705 |
| dist_settle:cos_ta | -37.750167 | 21.5796709 | -1.7493393 | 0.08023238 | -80.045545 | 4.54521075 | PYBI055 | model3 | 269.769705 |
| log_sl:cos_ta | 0.12824743 | 0.18528115 | 0.69217743 | 0.48882591 | -0.234897 | 0.49139182 | PYBI055 | model3 | 269.769705 |
| dist_road | 66.1428375 | 35.0596467 | 1.88658026 | 0.0592168 | -2.5728074 | 134.858482 | PYBI055 | model4 | 267.634621 |
| log_sl | 66.2305901 | 50.3810234 | 1.31459398 | 0.18864639 | -32.514401 | 164.975582 | PYBI055 | model4 | 267.634621 |
| cos_ta | 192.555114 | 119.755888 | 1.60789684 | 0.10785777 | -42.162114 | 427.272341 | PYBI055 | model4 | 267.634621 |
| dist_road:log_sl | -8.4362662 | 6.41095667 | -1.3159138 | 0.18820298 | -21.00151 | 4.12897795 | PYBI055 | model4 | 267.634621 |
| dist_road:cos_ta | -24.602038 | 15.2254994 | -1.6158444 | 0.10612796 | -54.443468 | 5.23939251 | PYBI055 | model4 | 267.634621 |
| log_sl:cos_ta | 0.13948193 | 0.18054502 | 0.77256037 | 0.4397826 | -0.2143798 | 0.49334367 | PYBI055 | model4 | 267.634621 |
| dist_water | 196.408748 | 133.314021 | 1.47327901 | 0.14067582 | -64.881931 | 457.699427 | PYBI055 | model5 | 258.010249 |
| log_sl | 110.670247 | 232.186039 | 0.47664471 | 0.63361514 | -344.40603 | 565.746521 | PYBI055 | model5 | 258.010249 |
| cos_ta | 485.696224 | 482.450009 | 1.0067286 | 0.31406521 | -459.88842 | Inf | PYBI055 | model5 | 258.010249 |
| dist_water:log_sl | -12.525599 | 26.3121062 | -0.4760394 | 0.63404631 | -64.09638 | 39.0451811 | PYBI055 | model5 | 258.010249 |
| dist_water:cos_ta | -55.142527 | 54.6582371 | -1.0088603 | 0.31304163 | -162.2707 | 51.9856496 | PYBI055 | model5 | 258.010249 |
| log_sl:cos_ta | 0.24183342 | 0.20061972 | 1.20543196 | 0.22803659 | -0.151374 | 0.63504085 | PYBI055 | model5 | 258.010249 |
| dist_aq.ag | 130.650766 | 72.0800944 | 1.81257762 | 0.06989699 | -10.623623 | 271.925155 | PYBI055 | model6 | 268.804548 |
| log_sl | 103.867681 | 117.898287 | 0.88099398 | 0.37832108 | -127.20871 | 334.944077 | PYBI055 | model6 | 268.804548 |
| cos_ta | 369.594919 | 274.35733 | 1.34712974 | 0.17793845 | -168.13557 | Inf | PYBI055 | model6 | 268.804548 |
| dist_aq.ag:log_sl | -10.860402 | 12.3234986 | -0.8812759 | 0.37816853 | -35.014015 | 13.2932116 | PYBI055 | model6 | 268.804548 |
| dist_aq.ag:cos_ta | -38.720434 | 28.6643872 | -1.3508202 | 0.17675302 | -94.901601 | 17.4607323 | PYBI055 | model6 | 268.804548 |
| log_sl:cos_ta | 0.21461585 | 0.18864062 | 1.13769694 | 0.25524705 | -0.155113 | 0.58434467 | PYBI055 | model6 | 268.804548 |
| dist_terr.ag | 28.6344599 | 84.9556207 | 0.33705198 | 0.73607772 | -137.8755 | 195.144417 | PYBI055 | model7 | 272.421182 |
| log_sl | -40.964677 | 146.062755 | -0.2804594 | 0.77912505 | -327.24242 | 245.313062 | PYBI055 | model7 | 272.421182 |
| cos_ta | 163.559284 | 337.12617 | 0.48515748 | 0.62756464 | -497.19587 | Inf | PYBI055 | model7 | 272.421182 |
| dist_terr.ag:log_sl | 4.96288186 | 17.7098622 | 0.28023266 | 0.77929901 | -29.74781 | 39.6735739 | PYBI055 | model7 | 272.421182 |
| dist_terr.ag:cos_ta | -19.956074 | 40.8790872 | -0.4881732 | 0.62542718 | -100.07761 | 60.1654648 | PYBI055 | model7 | 272.421182 |
| log_sl:cos_ta | 0.24816496 | 0.17528019 | 1.4158186 | 0.15682862 | -0.0953779 | 0.59170783 | PYBI055 | model7 | 272.421182 |
| dist_road | 14.4419514 | 12.3852079 | 1.16606451 | 0.24358837 | -9.8326101 | 38.7165129 | PYBI055 | model8 | 271.584402 |
| dist_forest | 7.25917516 | 5.32418806 | 1.36343327 | 0.172746 | -3.1760417 | 17.694392 | PYBI055 | model8 | 271.584402 |
| dist_settle | 12.454259 | 21.6497736 | 0.57526047 | 0.56511515 | -29.978518 | 54.8870356 | PYBI055 | model8 | 271.584402 |
| log_sl | -0.0329513 | 0.12812946 | -0.2571716 | 0.79704633 | -0.2840804 | 0.21817788 | PYBI055 | model8 | 271.584402 |
| cos_ta | -0.9635034 | 0.74982472 | -1.2849715 | 0.19880224 | -2.4331328 | 0.50612604 | PYBI055 | model8 | 271.584402 |
| log_sl:cos_ta | 0.21636178 | 0.17592373 | 1.22986125 | 0.21874907 | -0.1284424 | 0.56116595 | PYBI055 | model8 | 271.584402 |
| dist_road | 23.2067544 | 11.7387785 | 1.97693094 | 0.04804944 | 0.19917138 | 46.2143374 | PYBI055 | model9 | 252.337735 |
| dist_terr.ag | 58.7199749 | 33.0911006 | 1.77449447 | 0.0759814 | -6.1373904 | 123.57734 | PYBI055 | model9 | 252.337735 |
| dist_water | 117.059666 | 33.2396342 | 3.52168935 | 4.29E-04 | 51.9111798 | 182.208152 | PYBI055 | model9 | 252.337735 |

| | | | | | | | | | |
|---|---|---|---|---|---|---|---|---|---|
| log_sl | 0.18807245 | 0.15619405 | 1.20409482 | 0.22855293 | -0.1180623 | 0.49420717 | PYBI055 | model9 | 252.337735 |
| cos_ta | -1.2282692 | 0.8069363 | -1.522139 | 0.12797425 | -2.8098353 | 0.35329686 | PYBI055 | model9 | 252.337735 |
| log_sl:cos_ta | 0.34638243 | 0.19155647 | 1.80825231 | 0.07056724 | -0.0290614 | 0.7218262 | PYBI055 | model9 | 252.337735 |
| dist_water | 108.32889 | 34.2818676 | 3.15994715 | 0.00157798 | 41.1376638 | 175.520115 | PYBI055 | model10 | 258.7288 |
| dist_settle | 15.5305463 | 23.863534 | 0.65080663 | 0.51517132 | -31.241121 | 62.3022135 | PYBI055 | model10 | 258.7288 |
| dist_aq.ag | 11.0232775 | 31.2555182 | 0.3526826 | 0.7243264 | -50.236413 | 72.2829676 | PYBI055 | model10 | 258.7288 |
| log_sl | 0.14660393 | 0.15137333 | 0.96849242 | 0.3327985 | -0.1500824 | 0.44329021 | PYBI055 | model10 | 258.7288 |
| cos_ta | -1.1462094 | 0.81309141 | -1.4096931 | 0.15863031 | -2.7398393 | 0.44742051 | PYBI055 | model10 | 258.7288 |
| log_sl:cos_ta | 0.31746086 | 0.1951068 | 1.62711327 | 0.10371304 | -0.0649414 | 0.69986315 | PYBI055 | model10 | 258.7288 |

sl: step length, ta: turn angle, dist_* habitat feature (forest, settlement, road, water, aquatic agriculture, terrestrial agriculture).

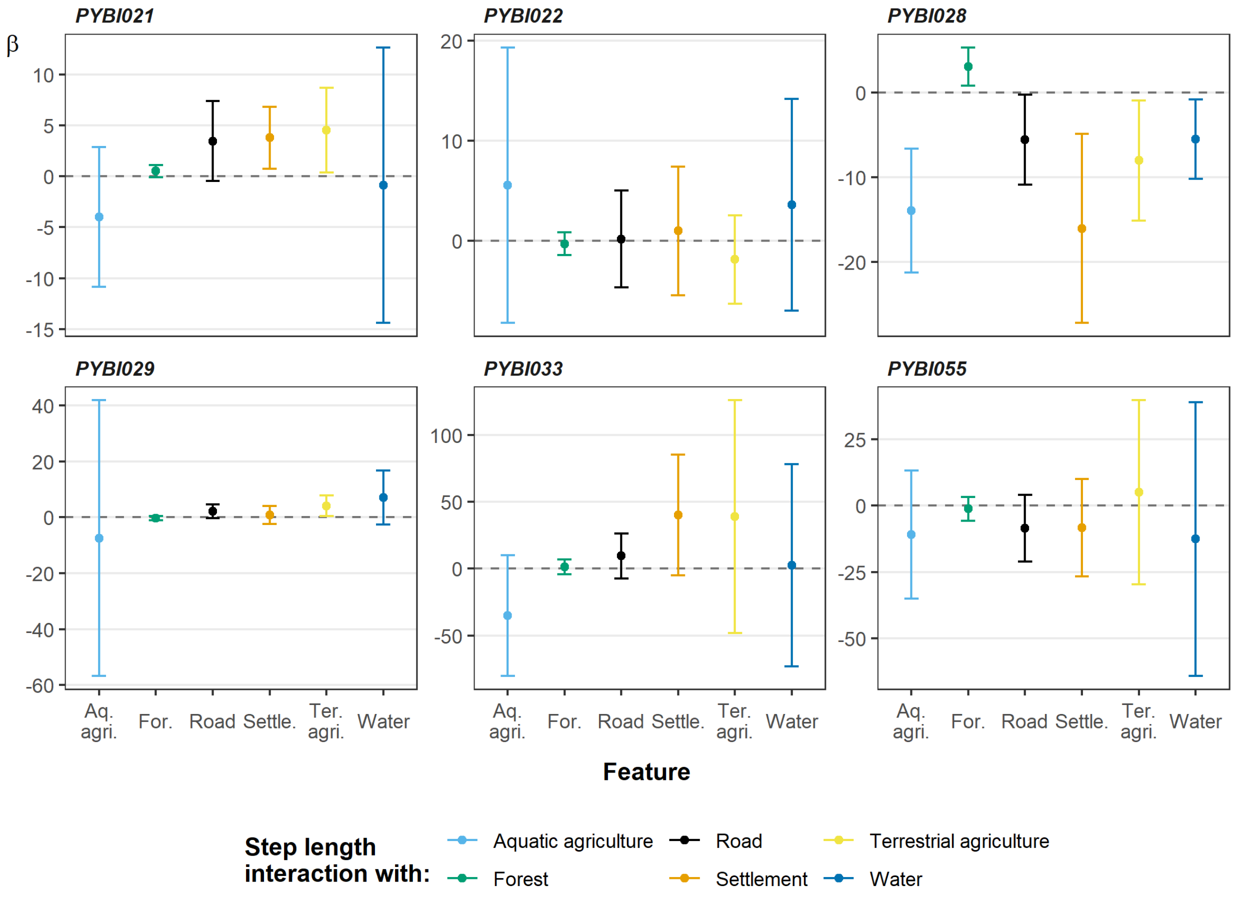


**Supplementary figure 2.** Interaction between step length (sl) and habitat feature for all individuals included in ISSF analysis. Error bars indicate 95% confidence intervals.

**Supplementary table 3.** All population level step selection analysis results.

| Mean | sd | q025 | q50 | q975 | Mode | kld | Term |
|---|---|---|---|---|---|---|---|
| 8.04E-04 | 7.16E-04 | -6.63E-04 | 8.11E-04 | 0.00222805 | 8.20E-04 | 8.36E-04 | dist_forest |
| 2.06E-06 | 1.35E-05 | -2.40E-05 | 1.93E-06 | 2.88E-05 | 1.69E-06 | 8.46E-08 | dist_forest:log_sl |
| -2.30E-05 | 2.66E-05 | -7.49E-05 | -2.30E-05 | 2.95E-05 | -2.32E-05 | 5.83E-07 | dist_forest:cos_ta |
| 5.66E-04 | 0.00109971 | -0.0012233 | 4.50E-04 | 0.00308549 | 3.18E-04 | 0.00340961 | dist_settle |
| 9.30E-08 | 4.09E-06 | -7.82E-06 | 5.77E-08 | 8.21E-06 | -1.21E-08 | 1.20E-07 | dist_settle:log_sl |
| -5.89E-06 | 7.98E-06 | -2.15E-05 | -5.91E-06 | 9.84E-06 | -5.96E-06 | 6.89E-07 | dist_settle:cos_ta |
| 0.0014194 | 0.00143973 | -0.0010807 | 0.0013038 | 0.00470286 | 0.0011849 | 5.27E-04 | dist_road |
| 1.32E-06 | 1.11E-05 | -2.01E-05 | 1.22E-06 | 2.33E-05 | 1.03E-06 | 7.20E-08 | dist_road:log_sl |
| -1.68E-05 | 2.16E-05 | -5.90E-05 | -1.68E-05 | 2.58E-05 | -1.70E-05 | 7.85E-07 | dist_road:cos_ta |
| 0.00272007 | 6.00E-04 | 0.00152945 | 0.00272433 | 0.00388592 | 0.00273284 | 3.54E-07 | dist_water |
| 8.18E-07 | 4.17E-06 | -7.27E-06 | 7.83E-07 | 9.10E-06 | 7.13E-07 | 2.10E-07 | dist_water:log_sl |
| -4.19E-06 | 8.16E-06 | -2.01E-05 | -4.21E-06 | 1.19E-05 | -4.26E-06 | 9.18E-07 | dist_water:cos_ta |
| -1.37E-04 | 7.73E-04 | -0.0016404 | -1.59E-04 | 0.00148567 | -1.98E-04 | 2.56E-05 | dist_aq.ag |
| -5.87E-08 | 1.97E-06 | -3.87E-06 | -7.55E-08 | 3.85E-06 | -1.09E-07 | 7.17E-08 | dist_aq.ag:log_sl |
| -2.52E-06 | 3.86E-06 | -1.01E-05 | -2.54E-06 | 5.08E-06 | -2.56E-06 | 8.96E-07 | dist_aq.ag:cos_ta |
| 0.00416689 | 0.00444432 | -4.77E-04 | 0.00241555 | 0.01611577 | 0.00165222 | 9.88E-05 | dist_terr.ag |
| 5.35E-07 | 7.44E-06 | -1.39E-05 | 4.71E-07 | 1.53E-05 | 3.45E-07 | 3.31E-07 | dist_terr.ag:log_sl |
| -9.22E-06 | 1.46E-05 | -3.77E-05 | -9.27E-06 | 1.95E-05 | -9.37E-06 | 5.32E-07 | dist_terr.ag:cos_ta |

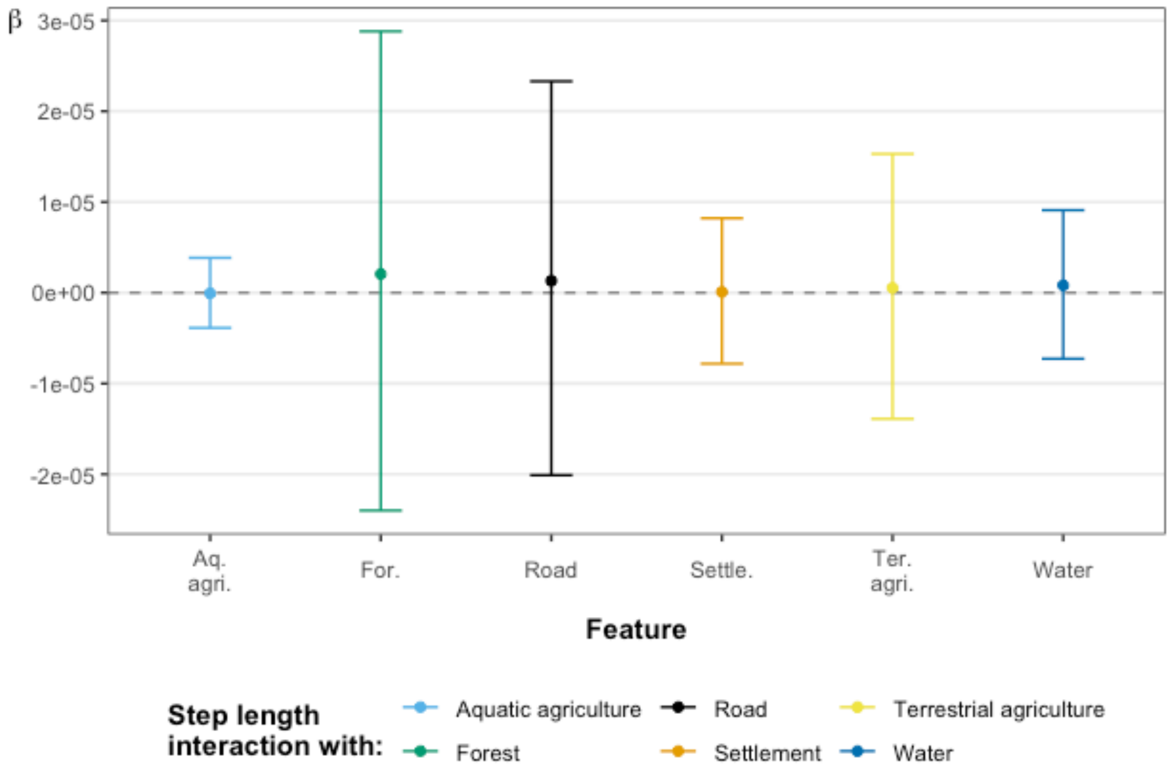


**Supplementary figure 3.** Interaction between step length and habitat feature at the population level. Error bars indicate 95% credible intervals.
